# RAB5c controls the assembly of non-canonical autophagy machinery to promote phagosome maturation and microbicidal function of macrophages

**DOI:** 10.1101/2025.03.25.645097

**Authors:** Edismauro Garcia Freitas-Filho, Isabella Zaidan, Marlon Fortes-Rocha, Daniel Leonardo Alzamora-Terrel, Carolina Bifano, Patrícia Alves de Castro, Renan Eugênio Araujo Piraine, Camila Figueiredo Pinzan, Caroline Patine de Rezende, Emilio Boada-Romero, Fausto Bruno dos Reis Almeida, Gustavo Henrique Goldman, Oliver Florey, Larissa Dias Cunha

## Abstract

Non-canonical conjugation of ATG8 proteins, including LC3, to single membranes implicates the autophagy machinery in cell functions unrelated to metabolic stress. One such pathway is LC3-associated phagocytosis (LAP), which aids in phagosome maturation and subsequent signaling upon cargo uptake mediated by certain innate immunity-associated receptors. Here, we show that a specific isoform of RAB5 GTPases, the molecular switches controlling early endosome traffic, is necessary for LAP. We demonstrate that RAB5c regulates phagosome recruitment and function of complexes required for phosphatidylinositol-3-phosphate [PI(3)P] and reactive oxygen species (ROS) generation by macrophages. RAB5c facilitates phagosome translocation of the V-ATPase transmembrane core, which is needed for ATG16L1 binding and consequent LC3 conjugation. RAB5c depletion impaired macrophage elimination of the fungal pathogen *Aspergillus fumigatus* and disruption of the V-ATPase-ATG16L1 axis increased susceptibility *in vivo*. Therefore, early endosome-to-phagosome traffic is differentially regulated to promote LAP and ROS contributes to resistance against *A. fumigatus* by effecting LAP.

**HIGHLIGHTS:**

- RAB5c is required for LC3-associated phagocytosis
- RAB5c finetunes NAPDH oxidase assembly and ROS generation in the phagosome
- RAB5c regulates V-ATPase assembly on the phagosome
- RAB5c and V-ATPase-ATG16L1 axis are required for the killing of *A. fumigatus*

## INTRODUCTION

Phagocytosis of invading microorganisms and dead cells plays critical roles in development, homeostasis, infection resistance, and immunoregulation^1–3^. The processes of phagosome formation and maturation are tightly controlled to ensure efficient cargo digestion and to initiate signaling cascades that modulate phagocyte function^4–6^. Consequently, defects in cargo recognition, engulfment, or degradation of phagosome content can lead to pathogenic outcomes.

Macroautophagy, hereafter referred to as autophagy, is a degradative pathway in which the formation of an autophagolysosomal compartment facilitates the clearance of damaged organelles, nutrient recycling, and removal of cytosolic pathogens^10^. Phagocytosis initiated by activation of transmembrane pattern recognition receptors (such as Toll-like receptors TLR2 and TLR4 and C-type lectin receptor Dectin-1), immunoglobulin receptor FCγR, and receptors for dying cells, triggers a non-canonical autophagy pathway known as LC3-associated phagocytosis (LAP)^3,7–9^. In contrast to autophagy, LAP utilizes components of the autophagy machinery to couple receptor activation with phagosome maturation, shaping phagocyte effector functions and its consequences upon gene expression. Deficiencies in LAP impair the clearance of diverse pathogens by macrophages, including *Aspergillus fumigatus*^5,11,12^, *Salmonella enterica*^13^, and *Listeria monocytogenes*^14–16^. The microbicidal activity of IFN-γ against *Toxoplasma gondii* also relies on LAP^17^. Moreover, LAP fine-tunes antigen processing and presentation by controlling phagolysosome maturation in dendritic cells^18,19^. Upon the uptake of cell corpses, LAP in myeloid cells sustains an immunosuppressive microenvironment that promotes tumor tolerance^8^.

LAP and autophagy are evolutionarily conserved yet distinct pathways. The ULK1/2-ATG13 complex that initiates autophagy is dispensable for LAP^8,20^. Instead, LAP requires the assembly of a unique Beclin-1-VPS34 class III phosphatidylinositol-3 kinase [PI(3)K] complex containing UVRAG and Rubicon (RUBCN) to generate phosphatidylinositol-3-phosphate [PI(3)P] on the phagosome membrane^3,21,22^. During LAP, but not autophagy, ionic imbalance caused by reactive oxygen species (ROS) production via NADPH oxidase 2 (NOX2) complex triggers the assembly of phagosome V-ATPase^20,23–25^. This process enables the tethering of the E3 ligase-like ATG16L1-ATG5-ATG12 complex to the phagosome, independently of autophagy components FIP200 and WIPI2b, and promotes the recruitment of the other components of the ATG8 lipidation machinery^7,8,26,27^. The covalent binding of ATG8 family proteins (MAP1LC3A/B/B2/C; GABARAP/L1/L2) to phosphatidylethanolamine (PE) and phosphatidylserine (PS) residues of the phagosome membrane controls phagolysosome maturation rate during LAP^11,14,20,28^.

Phagosomes are dynamic organelles that undergo continuous remodeling during maturation through sequential fusion with endosomes^4,29–31^. Membrane recognition and fusion require homotypic interaction between RAB-GTPases. Although RAB GTPase-mediated targeting and fusion of early endosomes (EE) and late endosomes (LE) with maturing phagosomes is well-documented^29–32^, the significance of inter-compartment communication for LAP is unknown. Meanwhile, despite the identification of several components of LAP^3^ how they are recruited to the phagosome and integrated into a unified signaling pathway remain poorly understood. Elucidating the spatiotemporal organization of LAP could provide critical insights for therapeutically targeting innate immune responses in the context of infections, cancer, and immune dysregulation.

Here, we investigated the importance of EE-phagosome interactions during LAP. We found that RAB5c, a specific isoform of EE GTPase RAB5, is required for LC3 lipidation on phagosomes. We show that RAB5c regulates the localization and activity of the class III PI(3)K complex and controls the traffic of the transmembrane core of the V-ATPase to the phagosome. We further reveal that the V-ATPase binding domain of ATG16L1 is required for resistance to infection in mice. This study establishes a differential role for RAB5c in controlling phagosome maturation during LAP and demonstrates that phagosome ROS exerts microbicidal activity against *A. fumigatus* by acting as a second messenger for V-ATPase assembly and phagosome maturation.

## RESULTS

### RAB5c depletion impairs LC3-associated phagocytosis

In mammalian cells, three paralogs of RAB5 GTPase - *Rab5a, Rab5b*, and *Rab5c* - are associated with the control of EE trafficking^4,33,34^, and descriptions of non-redundant functional roles between them are scarce^35–38^. Of note, the isoform RAB5c associates with phagosomes containing *Mycobacterium tuberculosis*^39^, *Salmonella enterica* serovar Typhimurium^40^, and *A. fumigatus*^12^, and is required for the maturation of the phagosome containing these pathogens.

Pathogenic *Legionella pneumophila* actively avoids fusion of RAB5c-containing EE with its vacuole through the activity of the bacterial effector VipD, arresting vacuole maturation and ultimately blocking lysosomal fusion^39,41,42^. This suggests that targeting RAB5c is a pathogen-driven subversion strategy to impair phagolysosome biogenesis, thereby allowing the establishment of a replicative niche. Real-time analysis of RAW264.7 macrophages expressing a mCherry-tagged murine RAB5c isoform (mCherry-RAB5c) showed the recruitment of mCherry-RAB5c to the vicinity of phagosomes containing opsonized zymozan (Op-Zym), a stimulus previously shown to induce LAP^7^. RAB5c recruitment occurred early upon Op-Zym uptake and lasted only during the first minutes of phagosome trafficking (**Video S1; Figs. S1A and 1A**). We also detected mCherry-RAB5c localized around phagosomes within the first 5 minutes of engulfment in macrophages stimulated with IgG-coupled Beads (IgG-Beads) to induce LAP but not in macrophages fed albumin-coupled Beads (BSA-Beads), which do not induce LAP and serve as an inert stimulus^20,25,43^. Our data indicate trafficking of RAB5c to early phagosomes during LAP (**Fig. 1B**).

**Figure 1.**
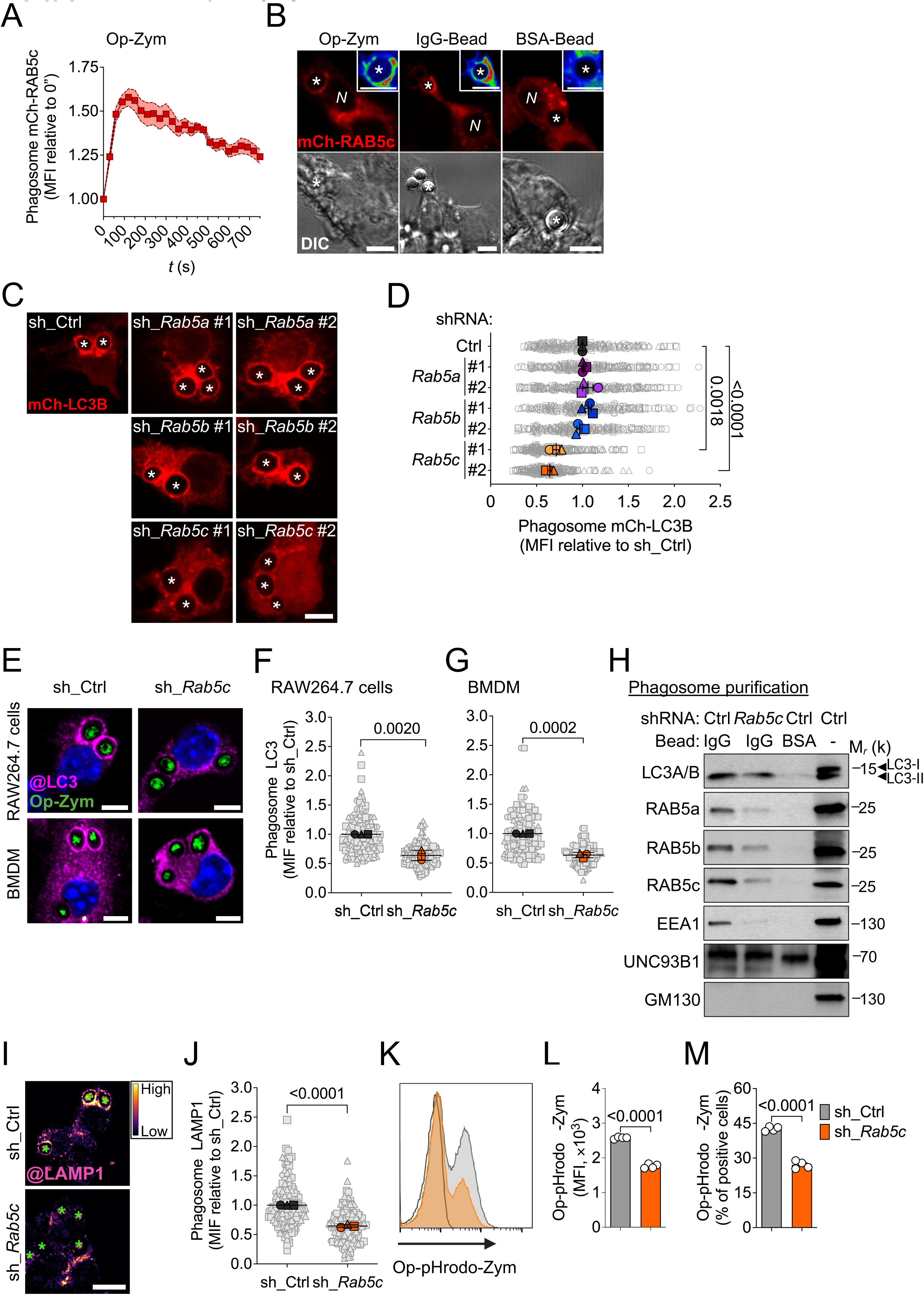
Rab5c depletion impairs LC3-associated phagocytosis. **(A)** Analysis of time-lapse confocal microscopy of mCherry-RAB5c (mCh-RAB5c) at phagosomes containing opsonized-zymosan (Op-Zym) in RAW264.7 cells. Fluorescence profile was normalized by the MFI at 0” for each phagosome. Squares are the mean and shaded area is ± S.E.M. for 12 phagosomes *(relative to Video S1)*. **(B)** Representative confocal images of RAW264.7-mCh-RAB5c cells stimulated for 5 min with Op-Zym, Beads adsorbed with IgG (IgG-Beads), or albumin (BSA-Beads). Insets show mCh-RAB5c fluorescence intensity using the RAINBOW LUT. *, phagosomes. N: nuclei. Scale bar: 5 μm. **(C-D)** Representative confocal images (C) and MFI quantification (D) of mCh-LC3B at phagosomes in RAW246.7 cells transduced with scrambled shRNA (sh_Ctrl) or indicated shRNAs (#1; #2) against *Rab5a, Rab5b, Rab5c,* and fed Op-Zym for 25 min. Scale bar: 5 μm. **(E-G)** Representative confocal images (E) and MFI quantification of LC3 immunolabeling (AF-647) at phagosomes in RAW264.7 cells (F) and BMDM (G) transduced with sh_Ctrl or sh_*Rab5c* and fed Op-Zym for 25 min. Blue: nuclei (Hoechst). Scale bar: 5 μm. **(H)** Immunoblot analysis of LC3, RAB5a/b/c and EEA1 in IgG-Bead- or BSA-Bead-containing phagosomes purified from RAW264.7 cells expressing sh_Ctrl or sh_*Rab5c* 40 min after stimulation. UNC93B1 was used as loading control and GM130, constitutive of Golgi apparatus, as control of purification. Data are representative of 4-5 experiments performed independently. **(I-J)** Representative confocal images (I) and MFI quantification (J) of LAMP1 immunolabeling (AF-488) at phagosomes in RAW264.7 cells fed Op-Zym for 25 min. The fluorescence intensity is shown using the FIRE LUT. Green *: phagosome. Scale bar: 10 μm. **(K-M)** Flow cytometric analysis of Op-pHrodo Red-Zym in RAW264.7 cells 45 min post-stimulation. Histogram (K), quantification of MFI (L), and percentage of positive cells (M) for each group are shown. The black line on the histogram represents the fluorescence of non-stimulated sh_Ctrl cells. Bars are the mean value of biological replicates (n=4, each indicated as a circle), and error bars are ± S.E.M. Statistical significance was calculated by unpaired Student′s *t*-test. Data shown is representative of 3 experiments performed independently. **(D, F, G, J)** Gray objects are the MIF of each analyzed phagosome. Colored objects are the mean value for each independent experiment (n=3) and error bars are ± S.E.M. Statistical significance was calculated by unpaired Student′s *t*-test between the indicated groups.

To probe the role of early endomembrane trafficking regulated by RAB5 during LAP, we performed gene silencing using two different shRNA constructs for each *Rab5* paralogue in RAW264.7 cells expressing fluorescent murine LC3B (mCherry-LC3B). We found that depletion of RAB5c, but not RAB5a or RAB5b, reduced the presence of mCherry-LC3B on phagosomes containing Op-Zym (**Fig. 1C and 1D**). We validated commercially available antibodies against each murine RAB5 isoform and confirmed silencing specificity (**Fig. S1B and S1C**). Real-time analysis confirmed that knockdown of RAB5c (*Rab5c* KD) impaired the conjugation of mCherry-LC3B to phagosomes instead of causing its removal **(Video S2; Fig. S1D**). Immunofluorescence confocal microscopy confirmed less endogenous LC3 decorating phagosomes containing Op-Zym in *Rab5c* KD RAW264.7 cells, without affecting particle uptake (**Fig 1E and 1F; Fig. S1E**). Similarly, *Rab5c* silencing in bone marrow-derived macrophages (BMDM) also reduced LC3 lipidation to phagosomes (**Fig. 1E-G, Fig. S1F and S1G**). Additionally, depletion of *Rab5c* by lentiviral CRISPR-Cas9 in RAW264.7 cells precluded LC3 recruitment to Op-Zym-containing phagosomes (**Fig. S1H-L)**, without affecting phagocytosis (**Fig. S1M)**.

Next, we performed phagosome purification in control and *Rab5c* KD RAW 264.7 macrophages. LC3-II was enriched in phagosomes containing IgG-Beads compared to BSA-Beads (**Fig. 1H; Fig. S1N**). Consistent with our microscopy observations, we found a reduction in LC3 levels in IgG-Bead-containing phagosomes from *Rab5c* KD macrophages (**Fig. 1H**). Notably, IgG-Beads-containing phagosomes from control macrophages were enriched in all RAB5 isoforms and the RAB5 effector EEA1 - a tethering factor that mediates endosomal vesicle docking and fusion with target membranes^30,44^ - compared to those containing BSA-Beads (**Fig. 1H; Fig. S1O-R)**. Conversely, *Rab5c* KD reduced EEA1, RAB5a, and RAB5b levels in the phagosomes with IgG-Beads, consistent with the idea that RAB5c facilitates early endosome recruitment to the nascent phagosomes during LAP (**Fig. 1H; Fig. S1O-R**).

Finally, we determined that RAB5c silencing reduced the levels of LAMP-1 in phagosomes containing Op-Zym, indicating that the fusion of late endosomes/lysosomes with phagosomes is impaired (**Fig.1J and 1K**). We next evaluated LAP-regulated acidification of phagosomes by feeding macrophages with Op-Zym labeled with the pH-sensitive probe pHrodo Red and performing flow cytometry. Phagosome acidification was reduced in *Rab5c* KD macrophages (**Fig. 1L and 1M**). These results confirm that RAB5c is required for phagolysosome maturation during LAP.

Collectively, these findings support that RAB5c regulates early endosome-phagosome communication during LAP and is required for LC3 conjugation and subsequent phagolysosome maturation.

### RAB5c depletion impairs PI(3)P production and stable assembly of NOX2 machinery on phagosomes but still increases phagosome ROS production

Next, we examined the role of RAB5c in the assembly of the LAP machinery onto the phagosome. First, we assessed the role of RAB5c on the recruitment and function of UVRAG-Beclin-1-VPS34-VPS15 class III PI(3)K complex during LAP. We found a reduction in VPS34, Beclin-1, and UVRAG levels on the phagosome membrane of IgG-Beads purified from *Rab5c*-silenced macrophages (**Fig. 2A; Fig. S2A-C**). RUBCN, which directly binds to VPS34 and is also required for PI(3)P generation on the phagosome membrane^26,45^, was also reduced on IgG-Bead-containing phagosomes from *Rab5c* KD macrophages (**Fig. 2A; Fig. S2D**). VPS34 is a RAB5 effector, and RAB5a recruits and induces VPS34 kinase activity in early endosomes^46^. To test the role of RAB5c in the function of VPS34 during LAP, we evaluate PI(3)P levels by imaging phagosomal association of a genetically-encoded probe based on the PI(3)P-binding domain of p40^phox^ (p40^phox^-PX-Venus)^47^. We found by real-time confocal analysis that the binding of p40^phox^-PX-Venus to phagosomes containing Op-Zym was reduced upon silencing of *Rab5c* (**Fig.2B and 2C; Video S3**). These findings corroborate that RAB5c helps recruit RUBCN and the class III PI3K complex onto phagosomes and is required for local PI(3)P generation during LAP.

**Figure 2.**
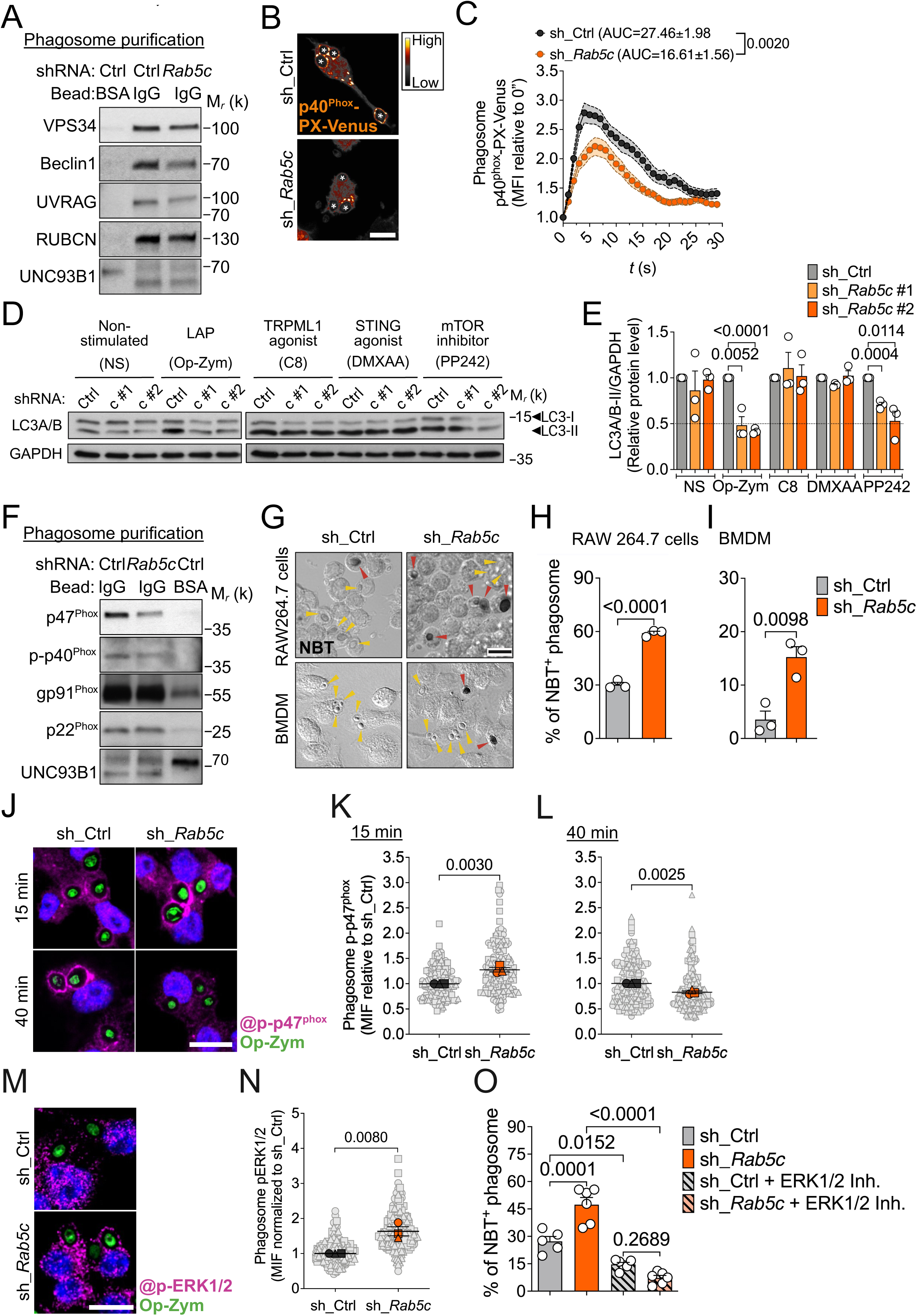
RAB5c depletion impairs PI(3)P production and stable assembly of NOX2 machinery on phagosomes but still increases phagosome ROS production. RAW264.7 cells or BMDM were transduced with scrambled shRNA (sh_Ctrl) or *Rab5c* shRNA (sh_*Rab5c*). **(A)** Immunoblot analysis of VPS34-PI(3)K CIII complex on BSA-Bead- or IgG-Bead-containing phagosomes purified from RAW264.7 cells at 40 min after stimulation. Data shown is representative of 3-5 experiments performed independently. **(B)** Representative confocal images of RAW264.7-p40^phox^-PX-Venus cells fed with Op-Zym for 25 min. The fluorescence intensity is displayed using the SMART LUT. *, phagosomes. Scale bar: 10 μm. **(C)** Quantification of time-lapse confocal microscopy analysis of p40^phox^-PX-Venus at phagosomes containing Op-Zym in RAW264.7 cells. Fluorescence profile was normalized by the MFI at 0” for each phagosome. Squares are the mean and shaded areas are ± S.E.M. for 18 phagosomes. The area under the curve (AUC) was determined for each analyzed phagosome and statistical significance was calculated by unpaired Student′s *t*-test. Data shown is representative of 2 independent experiments *(related to Video S3)*. **(D-E)** Immunoblot analysis (D) and densitometric quantification (E) of LC3A/B-II in RAW246.7 cells non-stimulated or incubated with Op-Zym for 25 min; C8 (2 μM) for 25 min; DMXAA (50 μg/mL) for 1 h; PP242 (1 μM) for 2 h, as indicated. GAPDH was used as loading control. Bars are the mean values of 3 independent experiments (each indicated as a circle), and error bars are ± S.E.M. Statistical significance was calculated by unpaired Student′s *t*-test. **(F)** Immunoblot analysis of NOX2 components on IgG-Bead or BSA-Bead containing phagosomes purified from RAW264.7 cells 40 min after stimulation. Data shown are representative of 4-5 experiments performed independently. **(G)** Representative DIC images of formazan deposits in phagosomes of RAW264.7 cells or BDMD fed Op-Zym for 15 min. Arrowheads indicate Op-Zym-containing phagosomes positive (Red) or negative (Yellow) for formazan deposition. **(H-I)** Percentage of NBT^+^ phagosomes containing Op-Zym in RAW264.7 cells (H) and BMDM (I). (**J-L)** Representative confocal images (J) and MIF quantification of p-p47^phox^ immunolabeling (AF-647) at phagosomes in RAW264.7 cells fed Op-Zym for 15 min (K) or 40 min (L).**(M-N)** Representative confocal images (M) and MIF quantification (N) of p-ERK1/2 immunolabeling (AF-647) in RAW264.7 cells fed with Op-Zym for 15 min. Blue: nuclei (DAPI). Scale bar: 10 μm. **(K, L, N)** Gray objects are the MIF of each analyzed phagosome. Colored objects are the mean value for each independent experiment (n=3) and error bars are ± S.E.M. Statistical significance was calculated unpaired Student′s *t*-test between the indicated groups. **(O)** Percentage of NBT^+^ phagosomes in RAW264.7 cells treated or not with ERK1/2 inhibitor (BVD-523; 100 μM) for 10 min and subsequently fed Op-Zym for 15 min. **(H, I, O)** Bars are the mean value of biological replicates (n=3-6, each indicated as a circle) and error bars are ± S.E.M. Statistical significance was calculated by unpaired Student′s *t*-test (H, I) or by one-way ANOVA and Tukey′s multiple comparisons (O) between the indicated groups. Data shown is representative of 3 (H, I) or 2 (O) experiments performed independently.

The class III PI(3)K complex containing UVRAG-Beclin-1-VPS34-VPS15 promotes autophagosome maturation by directing autophagosome fusion with late endosomes^48^, a function that does not require RUBCN^48–50^. Of note, we found that *Rab5c* depletion reduced autophagosome formation in response to macroautophagy induction with mTOR inhibitor PP242^51^, as assessed by immunoblot analysis of LC3-I to LC3-II conversion and the immunostaining of endogenous LC3 puncta by confocal microscopy (**Fig.2D and 2E; Fig. S2E**). Endolysosomal pH perturbations caused by multiple stimuli -such as viroporins, ionophores, lysosomotropic drugs, activation of endolysosomal channel TRPML1 (transient receptor potential cation channel, mucolipin subfamily, member 1), and STING (stimulator of interferon genes protein) activation - can also induce the non-canonical conjugation of LC3 to endolysosomal single membranes. In these cases, lipidation occurs independently of macroscale endocytic engulfment and RUBCN-VPS34 function^43^. Importantly, RAB5c depletion did not affect LC3-II levels in response to TRPML1 activation by the agonist C8 or activation of STING by DMXAA (**Fig. 2D and 2E; Fig. S2E**). These data further support that RAB5c is required for VPS34 kinase activity to induce lipidation of LC3 during LAP and is not universally required for non-canonical autophagy.

RUBCN and VPS34 contribute to the activity of the NADPH oxidase 2 (NOX2) complex in the phagosome membrane through the binding of transmembrane NOX2 p22^phox^ subunit to RUBCN^21^ and the cytosolic p40^phox^ subunit to PI(3)P, generated by VPS34^52^. NOX2 generates reactive oxygen species (ROS) within the phagosome lumen, establishing a RUBCN/PI(3)P/ROS axis essential for LAP progression^26,53^. We therefore examined the presence of NOX2 complex components onto phagosomes in the absence of RAB5c. IgG-Bead-containing phagosomes purified from *Rab5c* KD macrophages had reduced levels of cytosolic subunits p47^phox^ and phosphorylated p40^phox^ but retained similar levels of the transmembrane subunits p22^phox^ and gp91^phox^ when compared to phagosomes purified from control cells (**Fig. 2F; Fig. S2F-I**).

Next, we assessed phagosome ROS generation by measuring the deposition of insoluble formazan using the Nitroblue Tetrazolium (NBT) assay. Unexpectedly, we found an increased deposition of insoluble formazan mediated by O_2_- in Op-Zym-containing phagosomes from *Rab5c* KD RAW264.7 cells and BMDMs in comparison to control cells (**Fig. 2G-I**). We obtained similar results by measuring luminol oxidation during Op-Zym phagocytosis in the presence of Horseradish Peroxidase (HRP) (**Fig. S2J**). The elevated ROS burst in *Rab5c* KD macrophages in response to Op-Zym was suppressed by the NOX2 inhibitor diphenyleneiodonium (DPI) at nanomolar concentrations (**Fig. S2K**) and was not associated with increased levels of mitochondrial ROS (**Fig. S2L**), indicating that the higher ROS production remained NOX2-dependent. *Rab5c* depletion also increased ROS production in response to IgG-Beads (**Fig. S2M)** but not in response to Phorbol 12-Myristate 13-Acetate (PMA) **(Fig. S2N)**, which induces PKC-dependent NOX2 activation at the plasma membrane^54^. These results support that RAB5c modulates NOX2-mediated ROS production during LAP.

To understand how depletion of *Rab5c* could increase ROS levels despite reduced NOX2 complex assembly at the phagosome membrane, we imaged the translocation of cytosolic p47^phox^ - an early carrier of cytosolic p47^phox^ -p67^phox^-p40^phox^ ternary complex to phagosomes containing p22^phox55,56^ at different time points using confocal microscopy. In *Rab5c* KD macrophages, we found an early peak of p47^phox^ recruitment 15 minutes after Op-Zym internalization compared to control cells (**Fig. S2O, P)**. However, this increase was not sustained over time, with p47^phox^ levels declining at 40 minutes post-stimulation, consistent with our observations in purified phagosomes (**Fig. 2F**). Similarly, phagosomes from RAB5c-depleted macrophages exhibited a transient peak of decoration of p-p47^phox^, a phosphorylation event required for its binding to membrane-spanning p22^phox^ in murine cells^55^ (**Fig. 2J-L**). In human monocytes infected with *A. fumigatus*, active MAP kinase ERK1/2 traffics to the phagosome and controls p47^phox^ localization and phosphorylation, which is essential for ROS production during LAP^5^. Thus, we tested if ERK1/2 contributes to elevated ROS production in *Rab5c* KD macrophages. First, *Rab5c* silencing caused increased p-ERK1/2 decoration around Op-Zym-containing phagosomes (**Fig. 2M and 2N**). Second, pharmacological inhibition of ERK catalytic activity with Ulixertinib (BVD-523)^57^ abolished the enhanced ROS production in *Rab5c* KD cells, as assessed by the NBT assay (**Fig. 2O**). Collectively, these data suggest that RAB5c depletion hinders stable assembly of NOX2 on phagosome membrane. Nevertheless, the loss of RAB5c paradoxically increases phagosome ROS production by promoting ERK1/2 accumulation on phagosomes, which correlates with premature p47^phox^ activity.

### RAB5c controls the assembly of the V-ATPase in the phagosome during LAP

Because depletion of RAB5c did not cause a reduction of NOX2-mediated ROS production, we asked whether impairment of LC3 lipidation in *Rab5c* KD macrophages was due to regulation of other downstream events in the LAP cascade. During LAP, ROS-induced ionic imbalance triggers the binding of cytosolic V1 to the transmembrane V0 component of the phagosome V-ATPase^23,24^. ATG16L1 binds to ATP6V1H, a subunit of the V1 component of the V-ATPase, through its tryptophan-aspartic acid (WD) repeat-containing C-terminal domain^27^. This interaction enables the docking of the ATG12-ATG5-ATG16L1 complex onto single membranes and direct the site of LC3 lipidation^24,27,58,59^. Confocal microscopy analysis revealed a reduction in the engagement of ATP6V1A (a V1 subunit) on Op-Zym-containing phagosomes in *Rab5c* KD macrophages compared to control cells (**Fig. 3A and 3B**). Of note, the depletion of RAB5c did not affect the expression of ATP6V1A (**Fig. S3A and S3B)**. ATP6V1A levels were also reduced in IgG-Bead-containing phagosomes purified from *Rab5c* KD macrophages (**Fig. 3C; Fig. S3C**). Presence of ATG16L1 and ATG5/12 was also reduced on purified phagosomes of *Rab5c* KD macrophages, thus confirming that RAB5c is required for recruiting core ATG proteins that orchestrate the conjugation of LC3 to phospholipids (**Fig. 3C; Fig. S3D and S3E)**.

**Figure 3.**
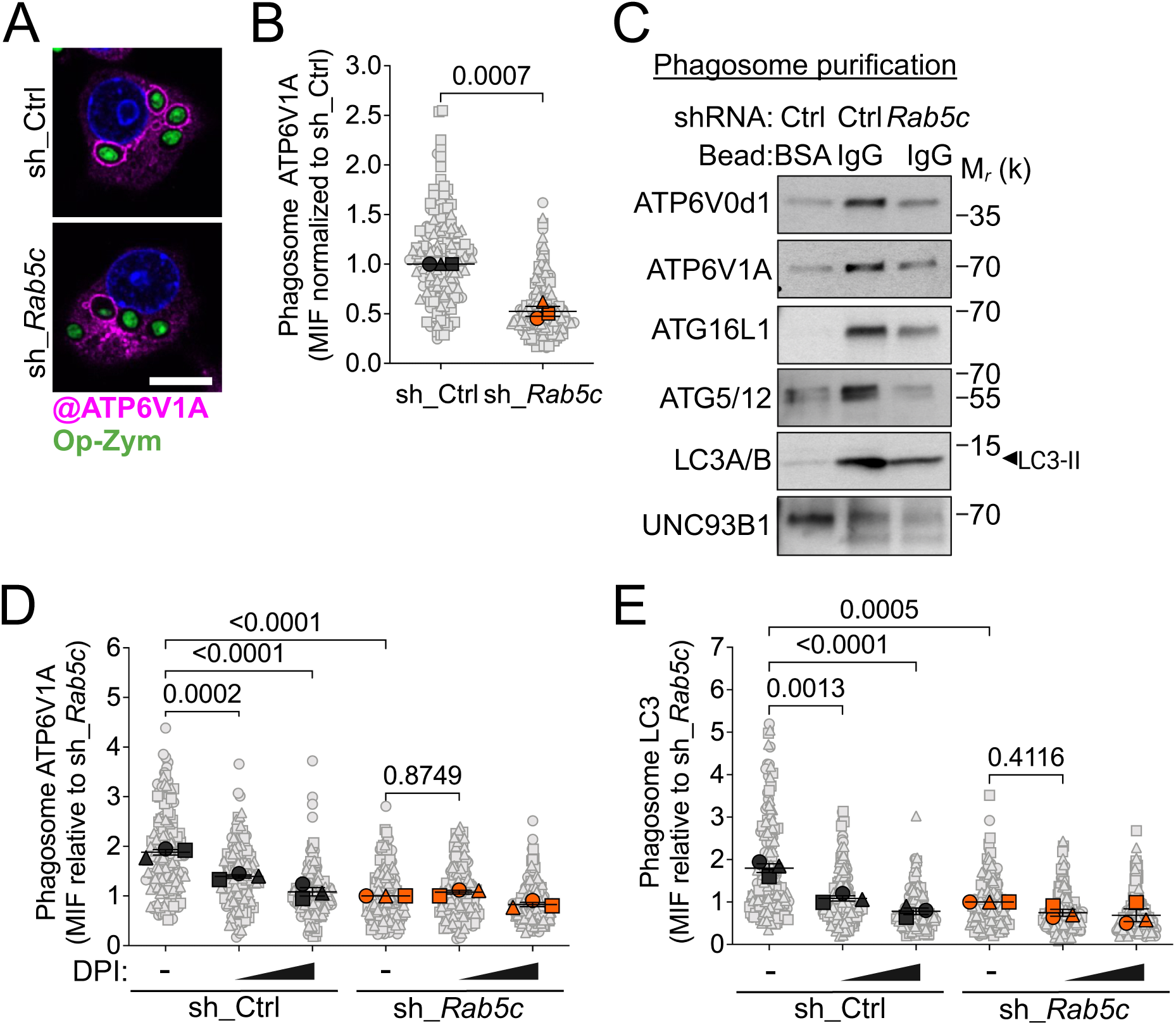
RAB5c controls the assembly of the V-ATPase in the phagosome during LAP. RAW264.7 cells were transduced with scrambled shRNA (sh_Ctrl) or *Rab5c* shRNA (sh_*Rab5c*). **(A, B)** Representative confocal images (A) and MFI quantification (B) of ATP6V1A immunolabeling (AF-647) at phagosomes in RAW264.7 cells fed Op-Zym for 25 min. Blue: nuclei (Hoechst). Scale bar: 10 μm. **(C)** Immunoblot analysis of components of the V-ATPase complex and ATG8 lipidation machinery on BSA-Bead- or IgG-Bead-containing phagosomes purified from RAW264.7 cells at 40 min after stimulation. Data are representative of 4-5 independent experiments. **(D-E)** MFI quantification of ATP6V1A (D) and LC3 (E) immunolabeling (AF-647) at phagosomes in RAW264.7 cells treated or not with DPI before feeding the cells with Op-Zym (lower concentration: 0.61 μM; higher concentration: 5 μM). **(B, D, E)** Gray objects are the MIF of each analyzed phagosome. Colored objects are the mean value for each independent experiment (n=3) and error bars are ± S.E.M. Statistical significance was calculated by unpaired Student′s *t*-test (B) or by one-way ANOVA and Tukey′s multiple comparisons (D, E) between the indicated groups.

Importantly, we found that RAB5c depletion decreased the levels of the V0 subunit ATP6V0d1 in purified IgG-Beads, indicating that RAB5c controls the traffic of the transmembrane core of the V-ATPase to the phagosome (**Fig. 3C, S3D**). This data could explain the impaired LC3 lipidation on phagosomes in RAB5c-deficient macrophages despite the presence of the triggering ROS signal. Alternatively, aberrant ROS production in *Rab5c* KD macrophages could disrupt V1 trafficking to the phagosome. To test the latter hypothesis, we treated *Rab5c* KD macrophages with DPI at low levels (61 nM) to normalize the ROS levels to those found in control cells (**Fig. S2K**). A high dose of DPI (5μM) abrogated ATP6V1A recruitment to the phagosome membrane in cells stimulated with Op-Zym (**Fig. 3D**). However, treatment with DPI at 61 nM failed to restore its recruitment in *Rab5c* KD cells (**Fig. 3D**). Consistently, ROS normalization in *Rab5c* KD macrophages did not rescue LC3 lipidation (**Fig. 3E**). Thus, lack of V-ATPase assembly upon RAB5c depletion is not directly cause by excessive ROS. Collectively, these findings support the notion that RAB5c participates in LAP by controlling the recruitment and activity of class III PI3K and the trafficking of the transmembrane V0 core of the V-ATPase to the phagosome.

### RAB5c-mediated V-ATPase assembly confers macrophage resistance to *Aspergillus fumigatus*

Generation of phagosomal ROS by NOX2 contributes for the elimination of *A. fumigatus* by macrophages^5,11,60,61^. Additionally, the fungal wall dihydroxy naphthalene (DHN)-melanin of *A. fumigatus* is a pathogenicity mechanism that excludes p22^phox^ of the phagosome and blocks LC3 lipidation^11^, further suggesting LAP activation is required for fungal killing and protection against invasive infection.

To investigate the importance of RAB5c and LAP for macrophage function, we first assessed its role in LAP induction upon engulfment of *A. fumigatus* conidia by BMDM using microscopy analysis. Silencing of *Rab5c* decreased LC3 conjugation to phagosomes containing *A. fumigatus* wild-type strain (ATCC 466645) (**Fig. 4A and 4B**). *Rab5c* silencing also impaired LC3 recruitment to phagosomes with the pigmentless melanin-deficient Δ*pksP* isogenic mutant, which induces LAP more efficiently^5,11,60^ (**Fig. 4A and 4B**). Using stimulation with Δ*pksP* conidia, we observed that reduced LC3 lipidation occurred despite increased ROS production in the phagosomes of *Rab5c* KD BMDM (**Fig. 4C and S4A**), mirroring the effect of LAP induction with Op-Zym (**Fig. 2G and 2I**). Of note, the increase in luminal ROS was associated with more recruitment of p-p47^phox^ and ERK1/2 to the phagosomes of *Rab5c* KD BMDM (**Fig. S4B and S4C**, respectively), thus corroborating that depletion of RAB5c disrupts normal ERK1/2 traffic and NOX2 function during LAP. Still, and consistent with our previous data (**Fig. 3A and 3B**), depletion of RAB5c resulted in less engagement of ATP6V1A onto phagosomes containing Δ*pksP* conidia (**Fig. 4D and S4D**), despite the increased ROS production.

**Figure 4.**
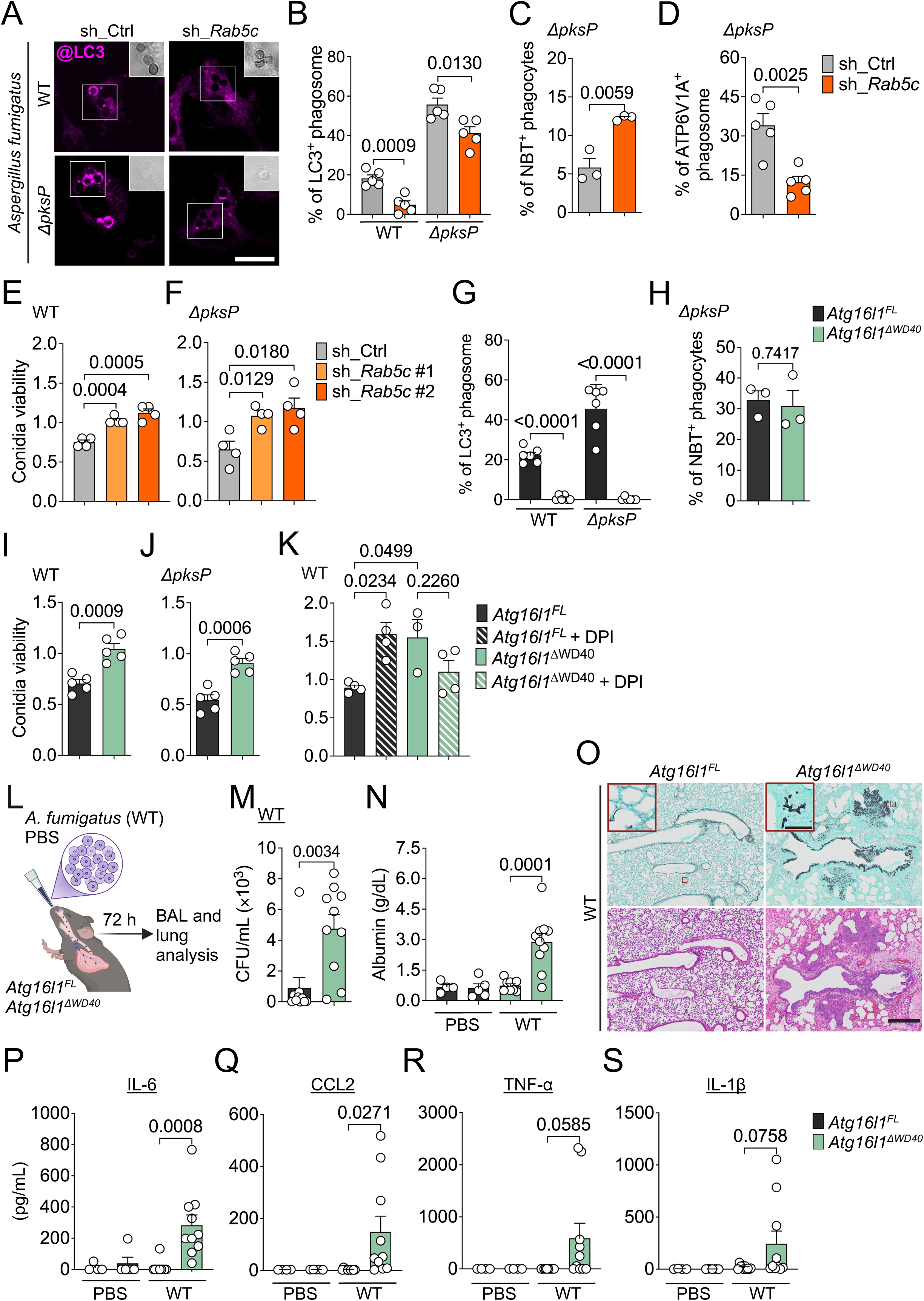
RAB5c-mediated V-ATPase assembly confers macrophage resistance to *Aspergillus fumigatus*. **(A-F)** BMDM transduced with scrambled shRNA (sh_Ctrl) or *Rab5c* shRNA (sh_*Rab5c*) were stimulated with conidia of wild-type (ATCC46645; WT) or Δ*pksP* strains of *A. fumigatus*. **(A, B)** Representative confocal images (A) and percentage of LC3-decorated phagosomes at 25 min of stimulation. Insets show details of conidia, scale bar: 10 μm. **(C)** Percentage of phagocytes positive for NBT deposition at 25 min of stimulation. **(D)** Percentage of ATP6V1A-decorated phagosomes at 1 h of stimulation. **(E, F)** Viability of WT (E) or Δ*pksP* (F) conidia determined as the ratio CFU at 6 h/CFU at 1 h. **(G-K)** *Atg16l1*^FL^ and *Atg16l1*^ΔWD40^ BMDM were stimulated with WT or Δ*pksP* conidia. **(G)** Percentage of LC3-decorated phagosomes at 25 min of stimulation. **(H)** Percentage of phagocytes positive for NBT deposition at 25 min of stimulation. **(I, J)** Viability of WT (I) or Δ*pksP* (J) conidia, as assessed by CFU counts as in (E). **(K)** Viability of WT conidia by BMDM pre-treated or not with DPI (5 μM), assessed by CFU counts as in (E). **(L)** Scheme of *A. fumigatus* lung infection. *Atg16l1*^FL^ and *Atg16l1*^*Δ*WD40^ mice were intranasally instilled with vehicle (PBS) or infected with WT *A. fumigatus* for 72 h. **(M)** Fungal loads in the bronchoalveolar lavage fluid (BAL) samples of infected mice, assessed by CFU counts. **(N)** Colorimetric quantification of BAL levels of Albumin. **(O**) Representative brightfield microscopy images of Grocott-Gomori′s methenamine silver (upper panel) or Hematoxylin and Eosin-stained (bottom panel) lung mice sections. Scale bar: 500 μm. Insets show details of *A. fumigatus* hyphae. Scale bar: 50 μm. **(P-S)** ELISA quantification of BAL levels of IL-6 (P), CCL2 (Q), TNF-α (R) and IL-1β (S). **(B-K)** Bars are the mean value of biological replicates (n=3-5, each indicated as a circle). **(M, N, P-S)** Bars represent the mean of 10 mice (each indicated as a circle). **(B-K, M, N, P-S)** Error bars are ± S.E.M. Statistical significance was calculated by unpaired Student′s *t*-test (B-J; M, N, P-S) or by one-way ANOVA and Tukey′s multiple comparisons (K) between indicated groups. Data shown are representative of 2 (A-D; G, H, K, M--S) or 3 (E, F, I, J) experiments performed independently.

Next, we addressed the importance of RAB5c for macrophage killing of *A. fumigatus.* We found that *Rab5c* silencing caused defective conidial elimination of wild-type (**Fig. 4E**) and Δ*pksP* (**Fig. 4F**) strains in comparison to control BMDM. Silencing of *Rab5c* did not affect the uptake of conidia (**Fig. S4E and S4F**). Collectively, our data support that RAB5c regulates LAP induction upon engulfment of *A. fumigatus* and is required for efficient clearance of the pathogen by macrophages.

Because RAB5c promotes V-ATPase assembly in the phagosome membranes, we sought to determine the mechanistic link between RAB5c, V-ATPase, and ATG16L1 to the resistance against *A. fumigatus*. To that end, we employed an *in vitro* infection model of BMDM from mice homozygously encoding two premature stop codons that prevent the translation of the ATG16L1 WD40 C-terminal domain (*Atg16l1^ΔWD40^*), which abrogates non-canonical lipidation of LC3 to single membranes without interfering with macroautophagy^25^. We found that conjugation of LC3 to phagosomes containing wild-type or Δ*pksP* conidia was severely impaired in *Atg16l1^ΔWD40^* BMDM (**Fig. 4G**). However, the expression of ATG16L1*^ΔWD40^* did not affect phagosome ROS production in response to infection, in agreement with NOX2 activation occurring earlier in the LAP cascade^15,21,24^ (**Fig. 4H**). *Atg16l1^ΔWD40^* BMDM internalized conidia similar to *Atg16l1^FL^*control cells (**Fig. S4G and S4H**) but eliminated wild-type and Δ*pksP* conidia less efficiently (**Fig. 4I and 4J**). Further, we found that treatment with DPI impaired the clearance of *A. fumigatus* in *Atg16l1^FL^* but not in *Atg16l1^ΔWD40^* BMDM, supporting that ROS generation by NOX2 requires V-ATPase and ATG16L1 axis to exert its microbicidal function (**Fig. 4K**). Similar to the expression of *Atg16l1^ΔWD40^* alleles, ablation of *Rubcn* did not affect the engulfment but impaired the killing of wild-type conidia (**Figs. S4I and S4J**, respectively), thus confirming the role of RUBCN in the control of *A. fumigatus* infection by macrophages.

Finally, we assessed the role of the V-ATPase-ATG16L1 axis in a nonlethal model of lung infection of immunocompetent mice with wild-type *A. fumigatus* conidia (**Fig. 4L**). Expression of ATG16L1^Δ*WD40*^ resulted in a higher recovery of viable conidia from the bronchoalveolar lavage fluid, suggesting impaired *in vivo* clearance of the pathogen by pulmonary phagocytes (**Fig. 4M**). Additionally, elevated albumin levels, indicative of lung damage, were detected in the lavage fluid of *Atg16l1^ΔWD40^,* but not *Atg16l1^FL^* mice, relative to their uninfected littermates (**Fig. 4N**). Histopathological examination revealed the presence of *A. fumigatus* hyphae within the lung parenchyma of *Atg16l1^ΔWD40^* mice 72 h post-infection (**Fig. 4O and Fig. S4K, S4L**). Further, lungs from infected *Atg16l1^ΔWD40^* mice presented clear signs of higher immune cell infiltration compared to *Atg16l1^FL^* mice (**Fig. 4O**). We also found increased levels of pro-inflammatory cytokines IL-6, CCL2, and TNF-α, and variable levels of IL-1β, in the lavage fluid of infected *Atg16l1^ΔWD40^* mice (**Fig. 4P-S**). Collectively, these data support the conclusion that disruption of the ATPase-ATG16L1 interaction results in impaired resistance and exacerbated immunopathology in response to *A. fumigatus*.

Altogether, our findings establish that LAP is required for efficient killing of *A. fumigatus* by macrophages and that non-canonical autophagy promotes immunity against infection *in vivo*.

## DISCUSSION

While RAB5 GTPases, markers of early endosome (EE) interactions, are frequently observed during the maturation of phagosomes formed upon recognition and internalization of pathogenic cargo, their functional role in this process remains poorly defined. Notably, RAB5 isoforms are often assumed to be functionally interchangeable, leaving gaps in our understanding of isoform-specific roles. Our findings demonstrate that RAB5c plays a non-redundant role in the LAP pathway during phagosome biogenesis, facilitating the efficient elimination of a fungal pathogens. Specifically, we show that RAB5c: 1) Orchestrates the transport and function of components responsible of PI(3)P generation, an early event of the LAP cascade; 2) Modulates the levels of ROS, a key second messenger; and 3) Enables LC3 lipidation by mediating the trafficking of the V-ATPase transmembrane core to the phagosome. By coordinating these processes, RAB5c ensures the spatiotemporal organization of the molecular platforms required for LAP signal transduction.

VPS34 is a known RAB5 effector, and RAB5a has been shown to recruit and activate VPS34 kinase function in endosomes^46^. At steady state, the RUBCN-UVRAG-Beclin1 complex localizes to EEA-1-positive endosomes independently of PI(3)P^62–65^. In this context, our data suggest that RAB5c may facilitate the transport of RUBCN-containing endosomes to phagosomes. However, whether RAB5c influences PI(3)P production by recruiting RUBCN or by directly binding to VPS34 on phagosomes remains to be determined. Notably, and similar to others^38,66^, we found that RAB5c silencing reduces LC3 lipidation during autophagy, a process that does not require RUBCN^7,10^. This observation suggests that RAB5c may independently regulate both the localization of RUBCN and its binding partners to phagosomes, as well as site-specific activation of VPS34.

Interestingly, recent evidence indicates that receptors triggering LAP share a common signature of phosphatidylserine (PS) clustering around a basic KRKPKK motif in their C-terminal domain, to which RUBCN can directly bind on the nascent phagosomes^67^. Only VPS34 that interacts with RUBCN can produce PI(3)P^67^. Therefore, it is possible that the assembly of a functional complex comprising RAB5c-VPS34-RUBCN-PS is required to stabilize PI(3)K CIII at the phagosome membrane once all components are recruited. Furthermore, it would be interesting to investigate whether PS clustering serves as the initial signal for RAB5c recruitment to phagosomes, given that K-Ras and RAC1 localization and activation are preferentially directed towards PS-rich membranes^68,69^.

The binding of p40^phox^ to PI(3)P is required for accumulation and stable retention of the ternary cytosolic NOX2 components at phagosomes^53^. In the absence of RAB5c, reduced PI(3)P generation may destabilize the NOX2 complex, potentially explaining the observed decrease in activated p40^phox^ and p47^phox^ in purified phagosomes following LAP induction. Notably, Ueyama *et al.* previously demonstrated that FcγR-mediated ROS production in macrophages depends on p40^phox^ PI(3)P binding only when p47^phox^ is partially phosphorylated at sites enabling its interaction with p22^phox^ ^52,55^. Thus, NOX2 activity and premature ROS burst in *Rab5c-*depleted macrophages may occur independent of prolonged NOX2 complex stability. In the absence of RAB5c on phagosomes, sustained ERK1/2 activity might drive oxidase activation, favored by p47^phox^. Since we confirmed previous findings that phosphorylated ERK1/2 (p-ERK1/2) localizes to phagosomes and is required for ROS production during *A. fumigatus* infection^5^, we speculate that RAB5c and EE interactions are necessary for the removal of p-ERK1/2 during phagosome maturation. This process could fine-tune ROS production, as excessive luminal ROS levels have been shown to accelerate V-ATPase assembly and LC3 lipidation on phagosomes while slowing acidification in monocytic cells^24^.

Our study demonstrates that RAB5c is required for the localization of V0 component of the V-ATPase to the phagosome, which subsequently enables V1 engagement in response to ROS. Our findings suggest that V0 recruitment is a regulated process dependent on communication between compartments of the endomembrane system. This idea aligns with previous biochemical evidence showing enrichment of the V0d1 subunit in EE^70,71^. Interestingly, real-time microscopy analysis revealed that the V0a3 component of V0 can be delivered to nascent phagosomes via tubular structures extending from lysosomes^72^. This observation suggests that phagosome-lysosome interactions may occur early in the maturation process. Further supporting this model, during LAP, localization of lysosomal marker LAMP-1 on phagosomes precedes LC3 lipidation^24^. Thus, while EE may not directly supply V0 to phagosomes, RAB5c-mediated phagosome maturation may facilitate early phagosome-lysosome interactions that cause V0 transfer. Additional studies will be necessary to determine whether RAB5c-regulated early endosome trafficking directly or indirectly governs the route delivering V0 subunits to phagosomes.

The discovery that ROS-dependent conjugation of LC3 to the phagosome membrane occurs in response to *A. fumigatus* infection led to the hypothesis that LAP, rather than a direct microbicidal mechanism, underlies the critical role of NOX2 and phagosomal ROS in macrophage resistance to the pathogen. Substantial data supporting this hypothesis comes from studies in murine models and human myeloid cells. However, the genetic basis for this connection has primarily relied on hematopoietic-specific ablation of *Atg5* or *Atg7*, genes shared by both LAP and autophagy, which renders macrophages and mice susceptible to infection^5,11,14,73,74^. The opposing effects of RAB5c on phagosomal ROS production and LC3 lipidation, by regulating different machineries that participate in LAP, allowed us to dissect the role of ROS in macrophage-mediated pathogen killing. Our findings demonstrate that NOX2-derived ROS eliminates *A. fumigatus* by triggering the V-ATPase-ATG16L1 axis of LAP. Macrophages deficient in RAB5C exhibited impaired V-ATPase assembly and LC3 lipidation, rendering them more susceptible to infection despite the elevated ROS levels. To provide definitive genetic evidence, we used a mutant mouse strain with disrupted V-ATPase-ATG16L1 interaction^25^, selectively impairing non-canonical autophagy. This model conclusively demonstrated that LAP is essential for macrophage-mediated *A. fumigatus* elimination. While our study establishes non-canonical autophagy as a critical mechanism for host resistance *in vivo*, future research is needed to determine the contributions of specific cell types to the susceptibility associated with *Atg16l1^ΔWD40^*.

## Supporting information

EGFF_etal_Supplemental File

## ACKNOWLEDGEMENTS

The authors thank Dr. Roberta R. C. Rosales, Msc. Elizabete R. Milani, Ms. Vani Maria Alves, Ms. Sandra Maria Oliveira Thomas, Ms. Rafaela Fernandes da Silva (Departamento de Biologia Celular e Molecular e Bioagentes Patogênicos, FMRP-USP, Ribeirão Preto, SP); Dr. Denise M. da Fonseca (Departamento de Imunologia, FMRP-USP); Dr. Patricia Vianna Bonini Palma (Hemocentro de Ribeirão Preto, Ribeirão Preto, SP); Dr. Simon Walker and Dr. Hanneke Okkenhaug (Babraham Institute, Cambridge, UK) for technical assistance.

## AUTHOR CONTRIBUTIONS

E.G.F.F., O.F. and L.D.C. designed the experiments. E.G.F.F. performed and analyzed the experiments. I.Z.M. and M.F.R. performed and analyzed specific experiments. D.L.A.T., C.B.A.A., P.A.C., R.E.A.P., C.F.P., C.P.R., and E.B.R. assisted with specific experiments. F.B.R.A., G.H.G., and O.F. provided resources. E.G.F.F. and L.D.C. wrote the manuscript and all the other authors revised it.

## MATERIALS AND METHODS

### Reagents

mTOR1 inhibitor PP242 (#4257) were purchased from Tocris; ERK1/2 inhibitor Ulixertinib (BVD-523; #HY-15816) from MedChemexpress. Hoechst 33342 Stain (H3570) was from ThermoFisher Scientific; Phorbol 12-myristate 13-acetate (PMA; tlrl-pma) was from InvivoGen; Bovine serum albumin (BSA; A4503); Sucrose (S7903); Imidazole (I2399); human IgG (I4506); human serum (P2918); NOX2 inhibitor: DPI (D2926); NBT (N6876); STING agonist, DMXAA (D5817); zymosan A (Z4250); pHrodo™ Red zymosan BioParticles™ Conjugate (P35364); MitoSOX™ Red mitochondrial superoxide indicator (M36008); horseradish peroxidase (HRP; P8375); Luminol (A8511); DAPI (D9542) were from Sigma Millipore Latex beads Polybead Microspheres 3.00 μm (17134-15) were from Polysciences. Murine IFNγ (315-05) was from Peprotech. TRPML1 agonist compound 8 (C8) was provided by Casma Therapeutics.

Compounds and drugs were reconstituted as per manufacturer′s recommendation and used as indicated in the text and figure legends.

### Antibodies

Anti-RAB5c [Thermo Fisher Scientific, PA5-39408; IB, 1:2,000; Immunofluorescence (IF), 1:300], anti-GAPDH [Cell Signalling Technology (CST), Clone D16H11, 5174T, IB, 1:10,000], anti-β-actin (CST, Clone 8H10D10, 12262, IB, 1:5,000), anti-Microtubule-associated protein 1 light chain B (anti-MAPLC3B/LC3B; CST, 27759, IB, 1:1,000), anti-LC3 (LC3A/LC3B/LC3C; MBL Int., PM036, IF: 1:250), anti-RAB5a (CST, 2143, IB, 1:1,000), anti-RAB5b (Santa Cruz, Clone F-9, sc373725, IB, 1:1,000), anti-EEA1 (CST, Clone C45B10, 3288, IB, 1:1,000), anti-UNC93B (Abcam, 69497, IB, 1:3,000), anti-GM130 (MBL, Clone 5G8, M1793, IB, 1:500), anti-UVRAG (CST, Clone D2Q1Z, 13115, IB, 1:1,000), anti-Phosphatidylinositol 3-kinase catalytic subunit type 3 (anti-PIK3C3/VPS34; Clone D9A5, CST 4263, IB, 1:1,000), anti-Beclin1 (CST, 3738, IB, 1:1,000), anti-Rubicon (Clone D9F7, CST, 8465, IB, 1:1,000), anti-p-p47^phox^(S328) (Abcam, 111855, IF, 1:50), anti-p47^phox^ (Abcam, Clone erp27205, 308256, IB, 1:1,000; IF, 1:250), anti-p-p40^phox^(Thr514) (CST, 4311, IB, 1:1,000), anti-p22^phox^ (Abcam, 75941 IB, 1:1,000), anti-gp91^phox^ (BD Bioscience, Clone53, 611414, IB, 1:2,000), anti-ERK1/2 (CST, Clone 1375F, 4695, IF, 1:500), anti-p-ERK1/2(Thr202/204) (CST, Clone 197G2, 9106, IF, 1:100), anti-ATP6V0d1 (Abcam, Clone erp18320-38, IB, 1:1,000), anti-ATP6V1A (Abcam, Clone erp19270, 199326, IB, 1:4,000; IF, 1:250), anti-ATG16L1 (MBL, Clone 1F12, M150-3, IB, 1:1,000), anti-ATG5/12 (CST, Clone D5F5U, 12994, IB, 1:1,000), anti-LAMP1 (CST, Clone E5N9Z, #99437, IF, 1:150), Alexa Fluor 647 polyclonal goat anti-rabbit IgG (Thermo Fisher Scientific, A21244, IF: 1:750), Alexa Fluor 488 polyclonal donkey anti-mouse IgG (Thermo Fisher Scientific, A21202, IF: 1:750), Alexa Fluor 647 polyclonal donkey anti-mouse IgG (Thermo Fisher Scientific, A31571, IF: 1:750), HRP-conjugated anti-rabbit IgG (CST, 7074, IB: 1:5,000), and HRP-conjugated anti-mouse IgG (CST, 7076, IB: 1:5,000) were used as indicated in manufactures′ datasheets.

### Aspergillus fumigatus culture

*A. fumigatus* (ATCC46645; WT) or *A. fumigatus* DHN-melanin-deficient strain (Δ*pksP*) were grown as described previously^75^ with slight modifications. Fresh *A. fumigatus* conidia were seeded on YAG medium supplemented with malt extract agar [2% (w/v) glucose, Sigma Millipore, G5767; 0.2% (w/v) yeast extract, Sigma Millipore, Y1625; 2% (w/v) malt extract, Gibco, 218630; 2% (w/v) agar, Oxoid, LP0011B] and 0.1% (v/v) trace element solution [ZnSO_4_, 221376 (76.5 mM); H_3_BO_3_, B0394 (178 mM); MnCl_2_, 244589 (40 mM); FeSO_4_, F8633 (18 mM); CoCl_2_, 449776 (12.3 mM); CuSO_4_, 451657 (10 mM); (NH_4_)_2_MoO_4_, 277908 (5.6 mM);

EDTA, 43178 (120 mM); all reagents were from Sigma Millipore] and cultured for five days at 37°C. Fungal culture for *in vivo* experiments was carried out without malt extract. Plates were washed with sterile PBS, followed by centrifugation and filtration through sterile Miracloth (Millipore, 475855) to obtain conidia suspensions. Conidia concentration was determined by light microscopy using a Neubauer chamber.

### Cells

HEK 293T (#CRL-3216) and Phoenix-Ampho cells (#CRL-3113) were obtained from ATCC. RAW264.7 macrophage cell line (ATCC, #SC-6003), HEK PEAKrapid cells (ATCC, #CRL-2828), and 3T3-MCSF cells were kindly provided by Dario Zamboni (Faculdade de Medicina de Ribeirão Preto). Cells were grown in monolayers in complete high glucose Dublbecco′s modified Eagle′s medium (DMEM; Gibco, 11995073) supplemented with contained 10% heat-inactivated fetal bovine serum [FBS (v/v), Gibco, 12657-029], 2 mM L-glutamine (Gibco, 35050061), and 100 IU/mL penicillin-streptomycin (Gibco, 15140122) (cDMEM media).

To prepare bone marrow-derived macrophages (BMDM), mice were euthanized, and bone marrow cells were harvested by flushing ethanol-sterilized femurs with DMEM media. BMDM differentiation was carried out by culturing cells for 6-7 days in cDMEM supplemented with conditioned medium of 3T3-MCSF cells [10%, (v/v)] and penicillin-streptomycin (100 U/mL). Differentiated BMDM were harvested by scraping in sterile phosphate-buffered saline (PBS) and seeded onto tissue culture plates in cDMEM media 24 h before stimulation.

All the cell lines were grown at 37°C in a humidified incubator with 5% CO_2_ in air and were periodically confirmed as mycoplasma negative by PCR according to^76^. All the reagents used for cell culture were purchased from Thermo Fisher Scientific.

### Mice

C57BL/6J (B6) mice are from Jackson Laboratories. *Atg16l1^ΔWD40^* mice^25^ were kindly provided by Thomas Wileman (Norwich Medical School, University of East Anglia, Norwich, Norfolk, UK). *Rubcn^-^*^/^- mice^8^ were kindly provided by Douglas Green (St. Jude Children′s Research Hospital, Memphis, TN, USA). *Atg16l1^ΔWD40^* and *Rubcn^-^*^/^- mice were previously crossed to C57BL/6 GFP-LC3^Tg^ mice (Noburo Mizushima, University of Tokyo, Tokyo, Japan). Mice were bred and housed in specific pathogen-free facilities at 23°C with a 12 h light/dark cycle in ventilated cages, with chow and water supply *ad libitum* at the Animal Research Facilities of the Faculdade de Medicina de Ribeirão Preto, Universidade de São Paulo, Ribeirão Preto, SP. Sex-matched mutant mice and background controls (8-12 weeks old) were used for experiments. The research was conducted in compliance with Ethical Principles in Animal Experimentation adopted by the National Council for Animal Experimentation Control. Experimental protocols were approved by the Ethics Committee on Animal Use of the Faculdade de Medicina de Ribeirão Preto (#1208/2023R1).

### Cell transfection and transduction

To construct RAW264.7 cell lines stably expressing RAB5 informs, a cDNA library generated from mouse placenta total mRNA (Takara, 636672) served as template to amplify the CDS mouse RAB5a (NM_025887.4), RAB5b (NM_177411.4) and RAB5c (NM_024456.5) sequences by PCR, using primers encoding 5′ MluI and 3′ NotI restrictions sites. pMX-mCherry-RAB5a, pMX-mCherry-RAB5b, and pMX-mCherry-RAB5c constructs were generated by cloning digested PCR products in-frame with a N-terminal mCherry fusion tag encoded in the MCS of a previously modified pMX-IRES-blasticidin vector (CellBiolabs #RTV-016)^67^. RAW264.7 cell lines stably expressing murine LC3B and p40^phox^-PX-Venus were constructed using pMXs-Venus-LC3B and pMXs-p40^phox^-PX-Venus that were previously described)^67^.

Phoenix-Ampho cells were used to produce Vesicular Stomatitis Virus-G (VSV-G) pseudotyped retrovirus to transduce RAW264.7 cells. Phoenix-Ampho cells were co-transfected using Polyethylenimine (PEI-MAX; 1 μg/mL) transfection reagent (Polysciences, 23966) and the pMX retroviral vector harboring the gene constructs of interest. Supernatants from Phoenix-Ampho cells were collected at 48 h post-transfection, filtered through 0.45 μm filters (Corning, 431220) and transduction was performed by spinfection (800×*g*; 40 min at 37°C). The medium containing the virus was removed and replaced with fresh complete DMEM media containing blasticidin S (Millipore Sigma, 15205; 5 μg/mL) 72 h post-transduction for selection of transformed cells. To validate the specificity of the commercial antibodies for each RAB5 isoform from *M. musculus*, HEK293T cells were transfect with pMXs-mCherry-RAB5a, pMXs-mCherry-RAB5b, and pMXs-mCherry-RAB5c to overexpress RAB5a, RA5b, and RAB5c proteins fused with mCherry using PEI-MAX and cells were harvested 72 h post-transfection.

For RAB5 silencing, pLKO.5-puromycin lentiviral constructs encoding shRNA sequences targeting the Rab5 isoforms and a control sequence were purchased from Millipore Sigma: Isoform a: sh_*Rab5a* #1: 5′-GCTGGTCAAGAACGGTATCAT-3′ (#TRCN0000100798) and sh_*Rab5a* #2: 5′-CAAGCAGCCATAGTTGTGTAT-3′ (#TRCN0000100799); Isoform b: sh_*Rab5b* #1: 5′-AGCCAGCCCTAGCATTGTTAT-3′ (#TRCN0000311442) and sh_*Rab5b* #2: 5′-GGAAGTCTAGCCTGGTGTTAC-3′ (#TRCN0000381840); Isoform c: sh_*Rab5c* #1: 5′- GCTAAGAAGCTTCCCAAGAAT-3′ (#TRCN0000100749) and sh_*Rab5c* #2: 5′- GCAATGAACGTGAATGAAATT-3′ (#TRCN0000100747); Control shRNA (sh_Ctrl): 5′- CCGGCAACAAGATGAAGAGCACCAACTCGAGTTGGTGCTCTTCATCTTGTTGTTTTT- 3′ (#SHC202).

To produce lentiviral particles, HEK PEAKrapid cells were transfected with VSV-G (pMD2.G; Addgene, #12259), PAX2 (Addgene, #12260), and pLKO.5 using PEI-MAX. For RAW264.7 cells, transduction with filtered lentiviral particles was carried out as described above. Cell media was replenished at 72 h post-transduction with fresh media containing puromycin (Millipore Sigma, P8833; 5 μg/mL) to select transformed cells. For BMDM transduction, non-adherent bone marrow cells were collected at day 2 of differentiation and mixed with the HEK PEAKrapid supernatants containing the lentiviral particles. Media was replenished 72 h post-transduction (day 5 of differentiation) with fresh warm media containing 5 μg/mL puromycin; selected BMDM were collected at day 7 and seeded for further experiments.

For the deletion of *Rab5c* in RAW264.7 cells by CRISPR-Cas9, guide sequences targeting the second exon of *Rab5c* from *M. musculus* (NM_024456.5) were designed using CRISPick design tool (https://portals.broadinstitute.org/gppx/crispick/public). Two sequences were selected (gRNA #1: 5′-CCACGCTCACTGGTACTACT-3′; gRNA #2: 5′-CCAGGAGAGCACAATTGGAG-3′) and cloned into the lentiCRISPR-v2GFP backbone (kind gift from Brett Stringer, Griffith University, Queensland, Australia; Addgene, #82416) digested with BsmBI (Thermo Fisher Scientific, ER0451). To produce lentiviral particles, HEK PEAKrapid cells were transfected using PEI-MAX with pMD2.G, pPAX2, and LentiCRISPR-v2GFP constructs. Transduction of lentiviral particles was carried out as described above. After 48 h of lentiviral transduction, a pool of GFP cells were selected by cell sorting (BD FACSAria™ Fusion Flow Cytometer; BD Bioscience) and further expanded for further experiments.

### Fluorescent microscopy

For fluorescent live-cell imaging, RAW264.7 cells (0.5-1.5×10^5^/plate) were seeded onto 35 mm glass-bottomed dish (P35G-1.5-14-C) in 2 mL of cDMEM media and cultivated overnight in the presence of IFN-γ (200 U/mL) prior to imaging. Opsonized-zymozan particles (Op-Zym) were generated by incubating zymosan A from *Saccharomyces* (1×10^6^ zymosan particle/μL) with human serum (v/v) for 30 min at 37°C. The solution was then centrifuged at 4,000×*g* for 5 min and resuspended in sterile PBS at 10 mg/mL. The solution was syringe-homogenised several times using a 26-G needle to break up aggregates. Plates were mounted on an Olympus SpinSR confocal microscope equipped with an Olympus IX83 stand, Olympus 60×1.5 NA UPLAPO objective lens, Yokogawa CSU-W1 scan head, and a Hamamatsu Orca Fusion camera. Images were acquired with a 2×2 camera bin and a 108 nmvpixel size. Laser excitation and emission filters for the mCherry and Venus channels were 561 nm (ex) 617/73 nm (em) and 488 nm (ex) 525/50 nm (em), respectively. The image acquisition software was Olympus cellSens v4.1. All imaging with live cells was performed within incubation chambers at 37°C and 5% CO2. For mCherry-RAB5c, p40-PX-dom-Venus, and mCherry-LC3B imaging, *z* stacks were acquired every 30 s following the addition of Op-Zym particles at a 1:4 macrophage:particle ratio. ImageJ software^77^ was used to measure the mean intensity of fluorescence in a region of interest (ROI) surrounding an individual Op-Zym particle and divided by the intensity of a ROI in the cytosol.

For fixed immunofluorescence confocal microscopy, 2.5×10^5^ RAW 264.7 cells or BMDM were seeded onto untreated 13 mm round coverslip in cDMEM (placed at the bottom of the well of a 24-well plate) and pre-incubated with IFN-γ (200 U/mL) for 16 h. Following treatments and stimulations, cells were rinsed twice with PBS and fixed for 20 min with 4% paraformaldehyde (w/v; Sigma Millipore, P6148) in PBS. Cells were quenched with 0.1 M glycine (Millipore Sigma, G7126) in PBS for 5 min. Samples that required immunostaining were permeabilized with 0.03% Triton X-100 (Millipore Sigma) for 10 min or with warm digitonin 200 mg/mL in PBS for 13 min. For immunolabeling of p-p47^phox^, p47^phox^, and ATP6V1A cells were fixed and permeabilized with 100% ice-cold methanol (Sigma Aldrich, 179337) for 5 min at −20°C. Next, cells were rinsed twice in PBS and incubated for 1 h at RT in PBS containing 1% BSA (w/v) and 5 μg/mL normal donkey IgG (Jackson ImmunoResearch Laboratories Inc., 017-000-003). Cells were labelled with primary antibodies diluted in PBS containing 1% BSA (w/v) for 1 h at RT or overnight at 4°C. Next, cells were rinsed thoroughly in PBS and incubated for 30 min at RT with the secondary antibodies diluted in PBS. For nuclear staining, after incubation with secondary antibodies, the cells were incubated for 5 min at RT with DAPI (0.2 μg/mL) or Hoechst 33342 (2 μM) in PBS. Cells were then rinsed in PBS, then in ddH_2_O, and mounted onto microscope slides with Fluoromount-G (Invitrogen, 00-4958-02) or with ProLong Gold Antifade Mountant (Invitrogen, P36934). Cells incubated without primary antibody served as negative controls. Samples were analyzed using a Zeiss LSM 780 laser scanning confocal microscope (Carl Zeiss Ltd) using Zen software (Carl Zeiss Ltd) or a LEICA STELLARIS 8 laser scanning confocal microscope (Leica Microsystems) using LAS X Life Science software (Leica Microsystems). High-resolution images were acquired using a Nikon Eclipse Ti2-E A1 high-resolution microscope (Nikon Instruments Inc.) using Nikon Elements software (Nikon Instruments Inc.). The MFI of the target protein on individual phagosomes were determined using ImageJ software. A mask drawn around the edge of a sampled phagosome and an adjacent 0.6 μm band were considered ROI to calculate the fluorescence intensity. For each condition, at least five images and forty phagosomes were quantified per experiment. For BMDM stimulated with *A. fumigatus,* phagosomes surrounded by a fluorescence rim of the indicated target protein were scored as positive. At least 50 phagosomes were analyzed for each condition per experiment, in a blinded fashion by the same investigator.

### Immunoblotting

1-3×10^6^ cells were washed twice with ice-cold PBS and immediately lysed with ice-cold RIPA buffer [Tris-HCl, pH 7.4, 50 mM; NaCl, 150 mM; EDTA, 1 mM; Triton X-100, T8787, 1% (w/v); SDS, L3771, 0.1% (w/v); Sodium deoxycholate, D6750, 0.5% (w/v); all reagents were from Sigma Millipore] supplemented with protease (cOmplete™; Roche, 11836153001) and phosphatase inhibitors (PhosSTOP™; Roche, 04906837001). Lysates were centrifuged at 16,000×*g* at 4°C for 20 min, and the supernatants collected. The protein content was quantified using the Pierce™ Bradford Protein Assay Kit (Millipore Sigma, 23200), with BSA as the standard^78^. Lysate containing 30 μg of protein was mixed with NuPAGE^®^ LDS Sample Buffer (Thermo Fisher Scientific) supplemented with 10% 2-betamercaptoethanol (v/v; Sigma Millipore, M3138) heated for 8 min at 90°C, and the proteins were separated electrophoretically by SDS-PAGE using Criterion XT Bis-Tris precast gels (4-12%; BioRad, 34501233/4/5) or 18% polyacrylamide (v/v; Sigma Millipore, A3574) SDS-PAGE gels and transferred to 0.45 μm polyvinydine difluoride membranes (Millipore, IPVH00010). Membranes were blocked for 1 h at RT in TBS-T [Tris-HCl, pH 7.5, 50 mM; NaCl, 150 mM; Tween 20, P9416, 0.05% (v/v)] supplemented with 4% BSA (w/v) and incubated overnight at 4°C or 1 h at RT with the individual primary antibodies diluted in blocking buffer. The membranes were then washed, incubated for 30 min at RT with the appropriate anti-IgG conjugated to HRP in TBS-T, washed and developed using SuperSignal™ West Pico Plus Chemiluminescent Substrate (Thermo Fisher Scientific) in an MI-5 X-Ray film processor (Medical Index) or with ChemiDoc XRS+ (Bio-Rad Laboratories, Inc.). After exposure, the films were digitized, and the mean optical density of the target protein was determined using ImageJ software. For stripping, after the first immunoblotting, the membrane was immersed in Restore™ Western Blot Stripping Buffer (Thermo Fisher Scientific, 21059) for 15 min at RT, washed, and the immunoblotting was repeated as described above.

### Phagosome purification

RAW264.7 cells were seeded onto 150 mm tissue-treated plate (Corning, 430599; 2×10^7^ cells/plate) in cDMEM and incubated for 16 h in the presence of IFN-γ (200 U/mL). Cells were fed IgG-Beads (human IgG coated latex Beads) or BSA-Beads (BSA coated latex Beads) at 1:8 macrophage:Bead ratio for 40 min and purified as previously described^67^. Briefly, plates were extensively rinsed with ice-cold PBS to eliminate non-internalized Beads. Cells were scraped in 10 mL of ice-cold PBS, pelleted, and mechanically homogenized in 1 mL 8.5% (w/v) sucrose (250 mM) through a 27-G needle. The cell homogenate was mixed with 62% (w/v) sucrose (181 mM) to reach a final sucrose concentration of ∼42%, layered onto 62% sucrose and carefully topped with 35% (w/v; 102 mM), 25% (w/v; 73 mM), and 10% (w/v; 29 mM) sucrose solutions in sequence in polycarbonate centrifuge tubes. Sucrose-containing buffers were prepared in 3 mM imidazole (pH 7.4). Following ultra-centrifugation at 100,000×*g* for 1 h at 4°C, phagosome-containing Beads were collected at the 10%-25% sucrose interface. Purified Bead-containing phagosomes were washed with ice-cold PBS, pelleted by ultra-centrifugation at 100,000×*g* for 20 min at 4°C, and resuspended in lysis buffer [Tris-HCl, pH 7.5, T6066; 50 mM; NaCl, 31434, 150 mM; EDTA, 5 mM; Igepal-CA-630, 13021, 1% (w/v); all reagents were from Sigma Millipore] supplemented with protease inhibitor (cOmplete™). Equal volume per sample was analyzed by SDS-PAGE and immunoblotting.

### Phagosome acidification assay

RAW264.7 cells were seeded onto 24 well plates (3.0×10^5^ cells/well) in cDMEM for 16 h in the presence of IFN-γ (200 U/mL). Cells were then fed with the pH sensitive probe pHrodo™ Red Op-Zym particles (1:2 macrophage:particle ratio) for 45 min. Cells were rinsed 3 times with ice-cold PBS, scraped, centrifuged at 300×*g* for 5 min, and resuspended in FACS buffer (BSA w/v, 1%, EDTA 1mM, in PBS). Sample acquisitions were performed in a FACSVerseTM (BD Biosciences) flow cytometer and data was analyzed with FlowJo™ v10.8 Software.

### ROS detection

For the NBT assay, RAW264.7 cells or BMDM were seeded onto untreated 13 mm round coverslip in cDMEM (placed at the bottom of the well of a 24-well plate) and cultivated in the presence of IFN-γ (200 U/mL) for 16 h. Prior to stimulation, cells were incubated with NBT in RPMI media (Gibco, 11835030) at 0.04 mg/mL for 10 min at 37°C. Where indicated, cells were also pre-incubated with ERK1/2 inhibitor prior to stimulation for 25 min. RAW 264.7 cells were stimulated with Op-Zym (1:8 macrophage:particle ratio) for 25 min and BMDM were stimulated with *A. fumigatus* melanin-deficient Δ*pksP* strain (1:4 macrophage:particle ratio) for 45 min. Samples were then fixed with 100% ice-cold methanol for 5 min at −20°C, extensively rinsed with PBS, once with ddH_2_O, and mounted onto microscope slides with Fluoromount-G. Slides were recorded by DIC or brightfield imaging on a Nikon Eclipse Ti2-E microscope (Nikon Instruments Inc.).

The luminol assay was performed as previously described^24^. Briefly, RAW264.7 cells (1×10^5^ cells/well) were seeded in cDMEM in 96-well white microwell plates (Thermo Fisher Scientific, 136101) and incubated in the presence of IFN-γ (200 U/mL) for 16 h. Cells were then rinsed with PBS and cell media replenished with sterile DPBS (Dulbecco′s PBS with Ca^2+^ and with Mg^2+^; Sigma Millipore, D8662) containing HRP (0.32 U/mL), luminol (125 nM), 0.1% glucose (m/v; Sigma Millipore, G8769), and sodium bicarbonate (4 mM, Sigma Millipore, S8761). Cells were incubated for 3 min at 37°C prior to stimulation. In some experiments, DPI were added to cell media 7 min before the incubation with the luminol/HRP solution and maintained until the end of the assay. Stimuli [Op-Zym, IgG-Beads, and BSA-Beads (1:8 macrophage:particle ratio) or PMA (500 ng/mL)] were added to the wells and measurements were obtained immediately in the MicroLumatPlus LB 96V (Berthold technologies) at 37°C.

To analyze mitochondrial ROS generation, RAW264.7 cells (3.0×10^5^ cells/well) were seeded on 24 well plates and incubated for 16 h in the presence of IFN-γ (200 U/mL). Cells were then washed twice with warm PBS and incubated with a pre-warmed MitoSOX Red mitochondrial superoxide indicator at 2.5 μM in serum-free DMEM media for 30 min prior to stimulation. Cells were fed Op-Zym (1:8 macrophage:particle ratio) for 30 min, rinsed with warm PBS, scraped from plates, centrifuged at 300×*g* for 5 min, and resuspended in FACS buffer. Cells were analyzed by flow cytometry (BD Biosciences, BD LSRFortessa Cell Analyzer) and data analysis was performed using FlowJo™ v10.8 Software.

### *Aspergillus fumigatus* killing assay

BMDM seeded in 96-well plates (1×10^5^ cells/well) were stimulated with *A. fumigatus* conidia at a 10:1 macrophage:conidia ratio. After 1 h incubation, cells were rinsed 3 times with warm sterile PBS to remove non-adherent, non-phagocytosed conidia, and media replenished with fresh cDMEM. Where indicated, 5 μM DPI was added 1 h before infection. At 1 h or 6 h post-infection, cells were lysed with sterile distilled H2O and cell lysate were seeded onto YAG-agar plates. CFU were counted after 24 h incubation at 37°C. Conidia viability was determined as the ratio CFU at 6 h/CFU at 1 h.

### *In vivo* infection with *Aspergillus fumigatus*

Mice were anesthetized with intraperitoneal injection of a solution of ketamine chloride (180 mg/kg) and xylazine chloride (20 mg/kg) and each challenged by intranasal instillation with vehicle or 1×10^9^ *A. fumigatus* conidia in 20 μL of PBS. At 72 h post-infection, mice were euthanized and the bronchoalveolar fluid (BAL) were collected in 1 mL of lavage with sterile PBS. 100 μL of the BAL was plated on YAG-agar for CFU counts. The remaining fluid was cleared by centrifugation and the supernatants were used for albumin quantification by a colorimetric assay (Labtest, #19-1/250) or IL-6 (DY406), CCL2 (DY479), TNF-α (DY410), and IL1-β (DY401) dosing by ELISA. All ELISA kits were acquired from R&D Systems and used according to the procedures supplied by the manufacturer. The lungs were harvested, then fixed in PBS with 4% PFA for 4 h and subsequently transferred to 70% ethanol for at least 24 h before paraffin embedding. 5 μm serial sections were stained with Hematoxylin and Eosin (HE) or Grocott-Gomori′s methenamine silver (GMS) and coverslips were mounted with Permount. Images were captured using an Olympus BX61 Motorized Slide Scanner Microscope Pred VS120 (Olympus, Hamburg, Germany).

### Data quantification and statistical analysis

Data were plotted and analyzed with GraphPad Prism version 10.3.0 (GraphPad Software, Boston, MA, USA; /https://www.graphpad.com). Please refer to the legend of the figures for description of samples (mice, cells or experimental replicates) and sample sizes. No statistical tests were used to estimate sample size. Statistical significance was calculated with either unpaired two-tailed Student′s t test or ANOVA, as specified in the legend of the figures. Paper figures were generated using BioRender.com. [Online] Available: https://biorender.com.

