## Supplementary material for "RAB5c controls the assembly of non-canonical autophagy machinery to promote phagosome maturation and microbicidal function of macrophages": EGFF_etal_Supplemental File

#### LEGENDS OF SUPPLEMENTAL FIGURES

**Supplemental Figure 1. Related to Figure 1.**

**(A)** Representative images of time-lapse confocal microscopy of mCh-RAB5c recruitment to a phagosome with Op-Zym (*) in RAW264.7-cells. Scale bar: 5 µm. **(B)** Immunoblot analysis of HEK293T cells, either wild-type or transfected with mCh-RAB5a, mCh-RAB5b, or mCh-RAB5c, using antibodies anti-RAB5a, anti-RAB5b, and anti-RAB5c. GAPDH was used as loading control. Red *: heterologous expressed RAB5 constructs. Arrows: endogenous RAB5 isoforms. Arrowheads: nonspecific bands. **(C)** Representatives immunoblot analysis of RAW264.7 cells transduced with sh_Ctrl or indicated shRNA sequences, using the validated antibodies in (B). GAPDH was used as loading control. **(D)** Quantification of mCh-LC3B MFI at Op-Zym-containing phagosomes of RAW264.7 cells transduced with sh_Ctrl or sh_*Rab5c*. Fluorescence profile was assessed by time-lapse confocal microscopy and normalized by the MFI at 0” for each phagosome. Squares are the mean and shaded areas are ± S.E.M. for at least 11 phagosomes. The area under the curve (AUC) was determined for each analyzed phagosome and statistical significance was calculated by unpaired Student's *t*-test. Data shown are representative of 2 independent experiments *(related to Video S2)*. **(E)** Percentage of Op-Zym phagocytosis by RAW264.7 cells expressing sh_Ctrl or sh_*Rab5c*, as assessed by immunofluorescence microscopy. **(F, G)** Immunoblot analysis (F) and densitometric quantification (G) of RAB5c levels in BMDM transduced with sh_Ctrl or sh_*Rab5c.* β-actin was used as loading control. **(H)** Scheme of the *Mus musculus Rab5c locus* with the identification of the CRISPR site for sgRNA#1 and sgRNA#2 in exon 2 (E2). **(I, J)** Immunoblot analysis (I) and densitometric quantification (J) of RAB5c in RAW264.7 cells non-transduced (NT), transduced with empty lentiviral vector backbone (EV), or *Rab5c* gRNAs. β-actin was used as loading control. **(K-M)** Representative confocal images (K), MFI quantification of LC3 immunolabeling (AF-647) at phagosome (L), and percentage of phagocytosis (M) in RAW264.7 cells transduced with EV or *Rab5c* gRNAs and fed Op-Zym for 25 min. Blue: nuclei (DAPI). Scale bar: 10 µm.

**(L)** Gray objects are the MIF of each analyzed phagosome. Colored objects are the mean value for each independent experiment (n=3) and error bars are ± S.E.M. Statistical significance was calculated by unpaired Student′s *t*-test between the indicated groups. **(E, G, J, M)** Bars are the mean values of 3 independent experiments (each indicated as a circle), and error bars are ± S.E.M. Statistical significance was calculated by unpaired Student′s *t*-test. **(N-R)** Densitometric quantitative analysis of LC3A/B (N), RAB5c (O), RAB5a (P), RAB5b (Q), and EEA1 (R) protein levels on purified phagosomes, using UNC93B1 as loading control. Bars are the mean value of 4-5 independent experiments (each indicated as a circle per condition with its connecting lines), and error bars are ± S.E.M. Statistical significance was calculated by unpaired Student′s *t*-test. (*related to Fig. 1H*).

**Supplemental Figure 2. Related to Figure 2.**

RAW264.7 cells or BMDM were transduced with scrambled shRNA (sh_Ctrl) or *Rab5c* shRNA (sh_*Rab5c*). **(A-D)** Densitometric quantitative analysis of VPS34 (A), Beclin1(B), UVRAG (C), and RUBCN (D) protein levels on purified phagosomes using UNC93B1 as loading control. Bars are the mean of 3-5 independent experiments (each indicated as a circle per condition with its connecting lines), and error bars are ± S.E.M. Statistical significance was calculated by unpaired Student′s *t*-test. (*related to Fig. 2A*). **(E)** Representative confocal images of LC3 immunolabeling (AF-647) in RAW264.7 cells non-stimulated (NS) or treated as in Fig. 3D. The fluorescence intensity is displayed using the ICA LUT. Scale bar: 10 μm. **(F-I)** Densitometric quantitative analysis of $p47^{phox}$ (F), p-$p40^{phox}$ (G), $gp91^{phox}$ (H), and $p22^{phox}$ (I) protein levels on purified phagosomes, using UNC93B1 as loading control. Bars are the mean value of 4-5 independent experiments (each indicated as circle per condition with its connecting lines), and error bars are ± S.E.M. Statistical significance was calculated by unpaired Student′s *t*-test. (*related to Fig. 2F*). **(J, K)** Assessment of luminol oxidation by relative quantification of chemiluminescence in RAW264.7 cells non-stimulated or fed Op-Zym in the presence of HRP.

In (K), cells were pre-treated or not with DPI at the indicated concentrations 10 min. Dots are the mean value of biological replicates (n=3), and error bars are ± S.E.M. Data shown is representative of 3 (J) or 2 (K) independent experiments. **(L)** Assessment of mtROS by MFI quantification of MitoSOX Red staining in RAW264.7 cells fed Op-Zym for 25 min. Bars are the mean values of 3 independent experiments (each indicated as a circle), and error bars are ± S.E.M. Statistical significance was calculated by unpaired Student's *t*-test. **(M-N)** Assessment of oxidative stress as in (J) in RAW264.7 cells stimulated with IgG-Beads or BSA-Beads (M), or PMA (500 μg/mL) (N). Dots are the mean value of biological replicates (n=3), and error bars are ± S.E.M. Data shown is representative of 3 independent experiments. **(O-P)** Representative confocal images (O) and MIF quantification (P) of $p47^{phox}$ immunolabeling (AF-647) at phagosome in RAW264.7 cells fed Op-Zym for 15 min or 40 min. Blue: nuclei (Hoechst). Scale bar: 10 μm.

**Supplemental Figure 3. Related to Figure 3.**

**(A, B)** Immunoblot analysis (A) and densitometric quantification (B) of ATP6V1A in the lysates of RAW264.7 cells transduced with sh_Ctrl or sh_*Rab5c*. GAPDH was used as loading control. Bars are the mean of 3 independent experiments (each indicated as a circle), and error bars are ± S.E.M. Statistical significance was calculated by unpaired Student's *t*-test. **(C-F)** Densitometric quantitative analysis of ATP6V0d1 (C), ATP6V1A (D), ATG16L1 (E), and ATG5/12 (F) protein levels on purified phagosomes, using UNC93B1 as loading control. Bars are the mean value of 4-5 independent experiments (each indicated as a circle per condition with its connecting lines), and error bars are ± S.E.M (related to Fig. 3C). Statistical significance was calculated by unpaired Student's *t*-test.

**Supplemental Figure 4. Related to Figure 4.**

**(A-F)** BMDM transduced with scrambled shRNA (sh_Ctrl) or *Rab5c* shRNA (sh_*Rab5c*) were stimulated with conidia of wild-type (ATCC46645; WT) or Δ*pksP* strains of *A. fumigatus*. **(A)** Representative DIC images of formazan deposits in phagosomes of BMDM stimulated with Δ*pksP* conidia for 45 min. Arrowheads indicate conidia-containing phagocytes positive (Red) or negative (Yellow) for formazan deposition, scale bar: 20 µm. **(B, C)** Percentage of p-p47$^{phox}$ (B) and ERK1/2-decorated (C) phagosomes at 25 min of stimulation. **(D)** Representative confocal images of ATP6V1A immunolabeling (AF-647) at phagosomes in BMDM infected with Δ*pksP* conidia. Scale bar: 10 µm. Blue: Calcofluor white (conidia). **(E, F)** Phagocytosis of WT (E) or Δ*pksP* (F) conidia at 1 h post-infection, assessed by CFU counts. **(G, H)** $Atg16l1^{FL}$ and $Atg16l1^{\Delta WD40}$ BMDM were infected with WT (G) or Δ*pksP* (H) conidia and phagocytosis was assessed by CFU counts. **(I, J)** $Rubcn^{+/+}$ and $Rubcn^{-/-}$ BMDM were infected with WT conidia and phagocytosis at 1h post-infection (I) and conidia viability at 6 h post-infection (J) was assessed by CFU counts. **(K)** Representative brightfield microscopy images of Hematoxylin and Eosin-stained lung mice, scale bar: 1,000 µm. **(L)** Representative brightfield microscopy images of Grocott-Gomori′s methenamine silver (left panel) or Hematoxylin and Eosin-stained (right panel) lung mice *(related to Fig. 4O)*, scale bar: 1,000 µm. **(B, C, E-J)** Bars are the mean value of biological replicates (n=4-6, each indicated as a circle), and error bars are ± S.E.M. Statistical significance was calculated by unpaired Student′s *t*-test between indicated groups. Data are representative of 2 (A-D, K, L) or 3 (E-J) independent experiments.

#### LEGENDS OF SUPPLEMENTAL VIDEOS

**Supplemental Video 1. Time-lapse spinning-disk confocal microscopy analysis of Op-Zym phagocytosis in RAW264.7 cells expressing mCherry-RAB5c.** Image stacks were acquired every 20 s. Movie plays 5 frames/s; time, min:s. mCh-RAB5c fluorescence intensity is scaled using the mpl-Plasma LUT. *Related to Fig. 1A*.

**Supplemental Video 2. Time-lapse spinning-disk confocal microscopy analysis of Op-Zym phagocytosis in RAW264.7 cells expressing mCherry-LC3B and transduced with sh_Ctrl or sh_*Rab5c*.** Image stacks were acquired every 30 s. Movie plays 3 frames/s; time, min:s. *Related to Fig. S1D*.

**Supplemental Video 3. Time-lapse spinning-disk confocal microscopy analysis of Op-Zym phagocytosis in RAW264.7 cells expressing p40$^{phox}$-PX-Venus and transduced with sh_Ctrl or sh_*Rab5c*.** Image stacks were acquired every 30 s. Movie plays 2 frames/s; time, min:s. mCh-RAB5c fluorescence intensity is scaled using the SMART LUT. *Related to Fig. 2C.*

### Freitas-Filho, et. al. 2025. Fig.S1

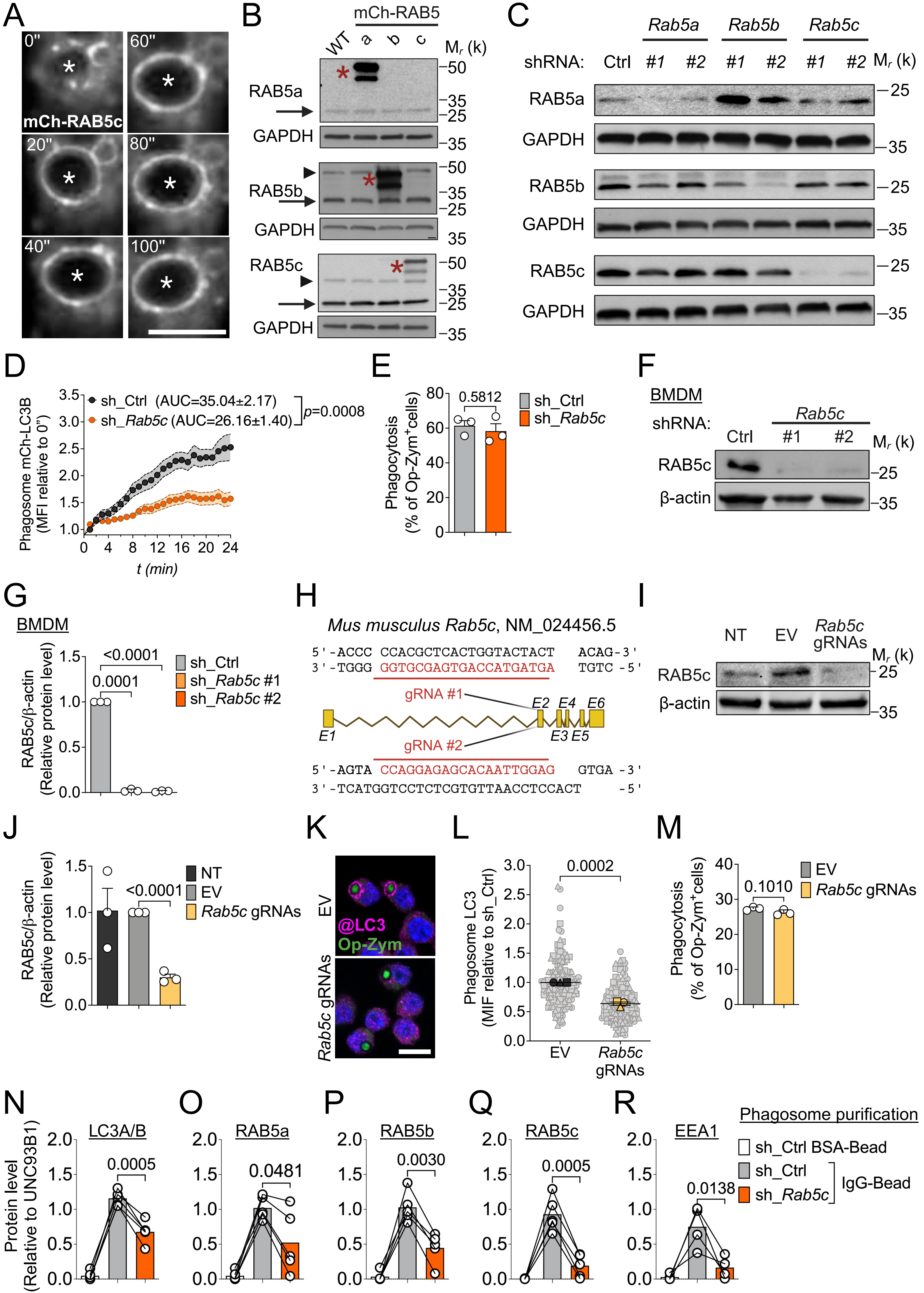

Freitas-Filho, et. al. 2025. Fig.S2

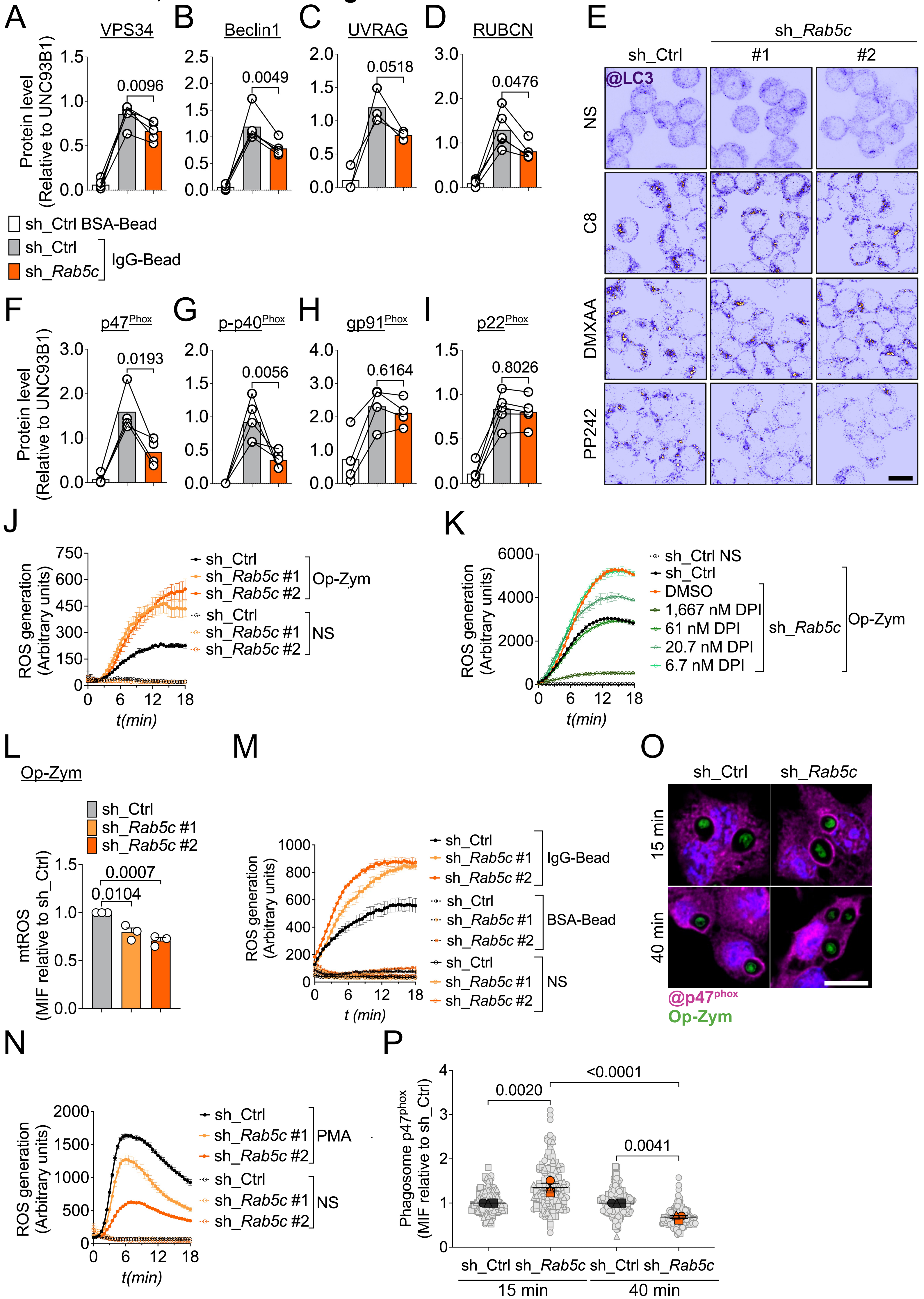

### Freitas-Filho, et. al. 2025. Fig.S3

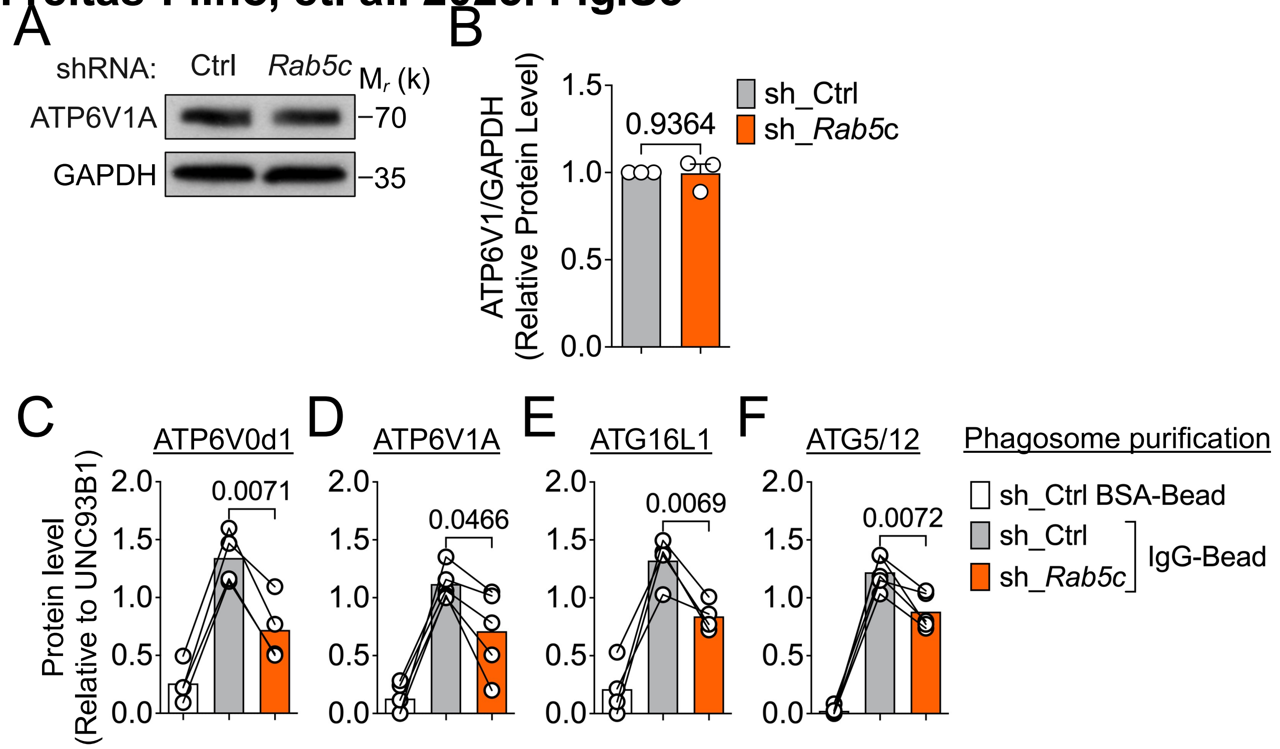

Freitas-Filho, et. al. 2025. Fig.S4

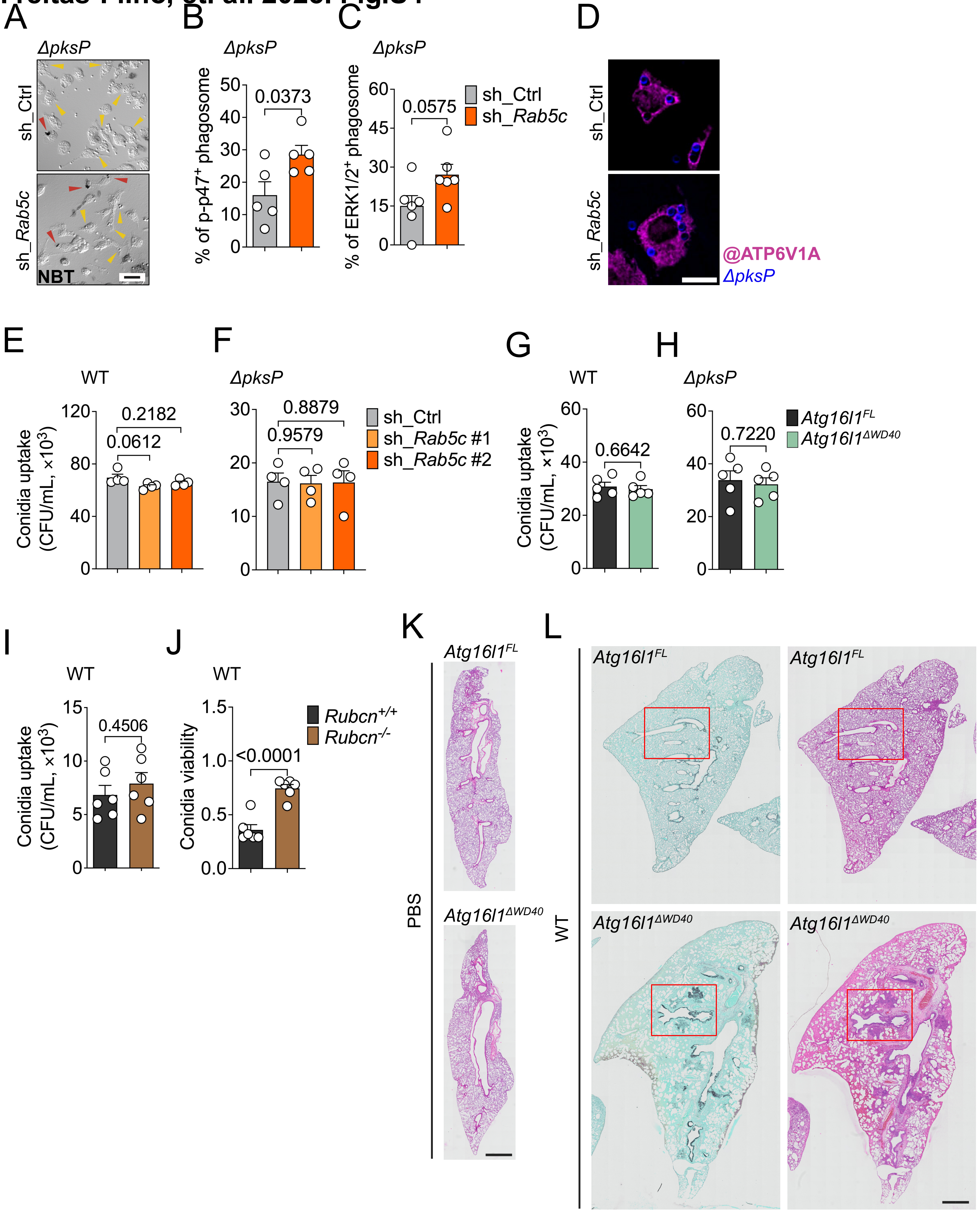
